# Negative effects of agricultural intensification on the food provisioning rate of a declining aerial insectivore

**DOI:** 10.1101/2021.03.24.436831

**Authors:** Daniel R. Garrett, Fanie Pelletier, Dany Garant, Marc Bélisle

## Abstract

The historical rise of intensive agricultural practices is hypothesized to be related to declines of grassland and aerial insectivorous birds. Drivers of declines may also influence the overall abundance and spatial distribution of insects within agricultural landscapes. Subsequently, the food provisioning rate of birds breeding within more agro-intensive landscapes may be impacted. Lower provisioning rates in agro-intensive landscapes may lead to reduced growth rate, body condition or fledging success of nestlings but also to diminished body condition of food provisioning adults. Results from a previous study supported this hypothesis as the fledging success and proxies of nestling body condition were lowest for an aerial insectivore breeding in more agro intensive landscapes. Of the multiple hypotheses put forward to explain these correlations, one mechanism may act through variation in food provisioning rates. In this study, we expounded on this hypothesis using data derived from the aforementioned study system and assessed if provisioning rates to nestlings and food provisioning behavior of adults varied across a gradient of agricultural intensification in a declining aerial insectivore, the Tree Swallow (*Tachycineta bicolor)*. We found that the hourly provisioning rate was lower in agro-intensive landscapes, and yet travel distances were longest within less agro-intensive landscapes. Our results highlight that, in order to maximize long term average gain rates, Tree Swallows breeding within agro-intensive landscapes must forage with greater intensity, perhaps at a cost to themselves, or else costs will transfer to growing broods. Our work provides further evidence that agricultural intensification on the breeding grounds can contribute to the declines of aerial insectivores in part through a trophic pathway.

## Introduction

Populations of both grassland and farmland birds have severely declined since the 1970s (Stanton et al. 2018, Rosenberg et al. 2019, Bowler et al. 2019). These groups include many aerial insectivores, a foraging guild presenting steeper population declines than most other avian guilds (Nebel et al. 2010, Michel et al. 2016). Concurrent changes in landscape composition and avian population indices have led to the hypothesis that these declines are driven, at least partly, by the increasing use of intensive agricultural practices (Benton et al. 2003, Hallmann et al. 2014, Stanton et al. 2018, Spiller and Dettmers 2019). Indeed, agricultural practices increasing agricultural yield per unit area of cultivated land, such as row cropping and the use of both synthetic fertilizers and pesticides, have risen dramatically in the past decades (Tscharntke et al. 2005, DiBartolomeis et al. 2019, Malaj et al. 2020).

Although many aerial insectivores share a common foraging strategy and diet, the ecological diversity of this guild invalidates many of the hypothesized drivers of farmland bird declines (Stanton et al. 2018, Spiller and Dettmers 2019). For instance, increased mechanization, including the frequency of forage harvesting, results in the significant nest destruction of ground nesting farmland birds (Tews et al. 2013, Stanton et al. 2018). Yet, most aerial insectivores do not nest on the ground and thus many hypothesized mechanisms underlying their declines involve a trophic pathway. One of those mechanisms postulates that intensive agricultural practices have reduced the overall abundance and changed the species composition or nutritive value of their prey (Stanton et al. 2018, Twining et al. 2018, Spiller and Dettmers 2019). Many factors of agricultural intensification can indeed affect the abundance, species assemblages, distributions, and trophic interactions of insects (Raven and Wagner 2021, Wagner et al. 2021). Therefore, intensive agricultural practices have the potential to affect the fitness of insectivorous birds through food limitation.

Previous studies have indeed observed that agricultural intensification, including habitats impactful to insect communities and abundances, correlate with changes in dietary composition (Nocera et al. 2012, English et al. 2017, Bellavance et al. 2018), reductions in annual fitness proxies (Benton et al. 2002, Ghilain and Bélisle 2008, Garrett et al. 2021a), and altered population growth (Stanton et al. 2018) of several aerial insectivore species. Studies linking prey availability to aerial insectivore breeding success have, however, often failed to observe such relationships (Dawson and Bortolotti 2000, Dunn et al. 2011, Imlay et al. 2017, McClenaghan et al. 2019). A lack of association between food availability and fitness proxies is potentially explained by breeding individuals compensating for reduced food availability or quality by altering foraging effort (Stephens et al. 2007). For altricial birds, food-provisioning is considered one of the most costly behaviors as individuals must meet the increasing metabolic demands of their brood while faring for themselves (Drent and Daan 1980, Wiersma 2005, Schifferli et al. 2014). Moreover, aerial insectivores are mostly short-lived species and thus life history theory suggests impediments to breeding will have significant impacts on lifetime reproductive success (Stearns 1992, Sæther and Bakke 2000, Berzins et al. 2020). Therefore, if compensation of reduced prey availability is through increased foraging effort, then it potentially comes at a cost to an individual’s own survival and future reproduction (Stearns 1992, Reznick et al. 2000, Harrison et al. 2011).

The large-scale homogenization of landscapes resulting from agricultural intensification (Benton et al. 2003) may thus alter foraging behavior and exacerbate the stress experienced by food-provisioning adults (Hinsley 2000, Olsson and Bolin 2014). Assuming that food-provisioning birds are maximizing their total daily food delivery, and if food patches are indeed of lower quality and/or more sparsely distributed within intensively cultivated landscapes, then central-place foraging theory predicts lower gain and nest visitation rates within these landscapes (Orians and Pearson 1979). Therefore, parents may optimize foraging strategies, potentially by increasing foraging efforts, to compensate for lowered food provisioning rates and overcome the consequences of nesting within such agro-intensive landscapes (Ydenberg 2007, Olsson et al. 2008). Empirical evidence for such elevated foraging workloads, often despite lower nest visitation rates, includes foraging at greater distances and reduced nest attendance in landscapes dominated by agriculture (Bruun and Smith 2003, Poulin et al. 2010, Catry et al. 2014, Stanton et al. 2016, Staggenborg et al. 2017, Evens et al. 2018). Furthermore, birds whose central places are located within more intensely cultivated landscapes present several conditions potentially related to increased foraging costs (Evens et al. 2018), including greater oxidative stress (Stanton et al. 2017, Evens et al. 2018) and lower immunocompetency (Belant et al. 2000, Pigeon et al. 2013). It is therefore imperative to assess the influence of landscape context on the capacity of breeding individuals to compensate for poor foraging conditions and the resultant workload.

A recent 11-year study in southern Québec, Canada suggested that landscapes dominated by more arable row crop monocultures (e.g. corn (*Zea mays*) or soybean (*Glycine max*)) offer significantly less aerial insectivore prey than ones dominated by less agriculturally intensive habitats such as forage crops, grasslands, and dairy farms (Garrett et al. 2021a). Furthermore, in this same study, and upon controlling for local prey availability, several indices of Tree Swallow (*Tachycineta bicolor*) breeding success remained associated with an agricultural gradient (Garrett et al. 2021a). Breeding attempts within landscapes dominated by intensively managed row crops resulted in reduced fledging success, longer nestling periods, and shorter fledging wing lengths. If these agro-intensive habitats do indeed reduce the functional connectivity of landscapes by reducing the amount and distribution of suitable foraging patches, then provisioning central-place foragers in these landscapes potentially expend more time or energy traveling between food patches and exploiting them, with the additional cost of having to settle for patches of lower quality (Hinsley 2000, Bélisle 2005, Olsson and Bolin 2014, Evens et al. 2018). Moreover, under such conditions, provisioning individuals may need to devote more time to self-feeding in order to maintain or increase their workload (Martins and Wright 1993, Markman et al. 2002, Ydenberg 2007). Therefore, one hypothesized mechanism explaining the results of Garrett et al. (2021a) is that swallows breeding in agro-intensive landscapes provide food at lower rates with the consequence that nestlings experience slower growth and potentially increased mortality.

Presented here are the results of a study evaluating the strength of evidence for this hypothesis using the same study system as Garrett et al. (2021a). Our main objective was to evaluate how a gradient of agricultural intensification influenced the provisioning rate and travel distances of food provisioning female Tree Swallows. Specifically, we evaluated a series of hypotheses grounded in central-place foraging theory (Olsson et al. 2008, Olsson and Bolin 2014). Provisioning models can address the constraints and mechanisms underlying the food provisioning behavior of Tree Swallows, notably because they take into account the time parents may devote to self-maintenance (Ydenberg 2007). However, a simple central-place foraging model, such as the marginal value theorem, likely provides the most parsimonious conceptual framework to make sensible qualitative predictions about provisioning rates. We hence firstly assumed the duration of each foraging bout is composed of the travel to, between, and within foraging patches, and that the biomass of food collected while exploiting a patch increases, yet at a decreasing rate with patch residency time (Olsson et al. 2008). Despite the uncertainty around which fitness currency provisioning altricial birds attempt to maximize, the fact that predation risk is likely negligible for adult foraging swallows and that agricultural intensification can reduce prey availability, if they are to maximize total daily food delivery, provisioning swallows may mostly be constrained by time, rather than physiology (Schoener 1971, McNamara and Houston 1997, Ydenberg 2007, Houston and McNamara 2014). We therefore further assumed food bolus biomass is positively related to its energy content, that traveling and foraging costs are constant per time unit, and that nestling rearing females are attempting to maximize their average long-term net rate of energy gains per time unit. Therefore, females whose nests are within landscapes composed of more dispersed or poorer food patches, as expected in landscapes dominated by agro-intensive cultures, will experience longer travel times (Appendix S1: Fig. S1A and C) or lower instantaneous gain rates while exploiting patches (Appendix S1: Fig. S1B and C), and thereby show lower food provisioning rates. Finally, shorter travel times can counterbalance slower gain rates within food patches, resulting in different food provisioning rates and yet similar net gain rates between contrasting landscapes (Appendix S1: Fig. 1D). Central-place foragers should be willing to incur greater time costs to reach and exploit food patches of a given quality in landscapes composed of more dispersed or poorer food patches (Olsson and Bolin 2014). We therefore tested if foraging distances would be furthest within more agro-intensive landscapes, indicating greater travel time and costs in such landscapes. We also tested the expectation that these effects would be at least partly alleviated with increasing local prey availability (i.e. prey availability close to the nest), but exacerbated by increasing food-demand from the brood (e.g., larger and older broods; Martins and Wright 1993, Markman et al. 2002, Geary et al. 2020).

**Figure 1:**
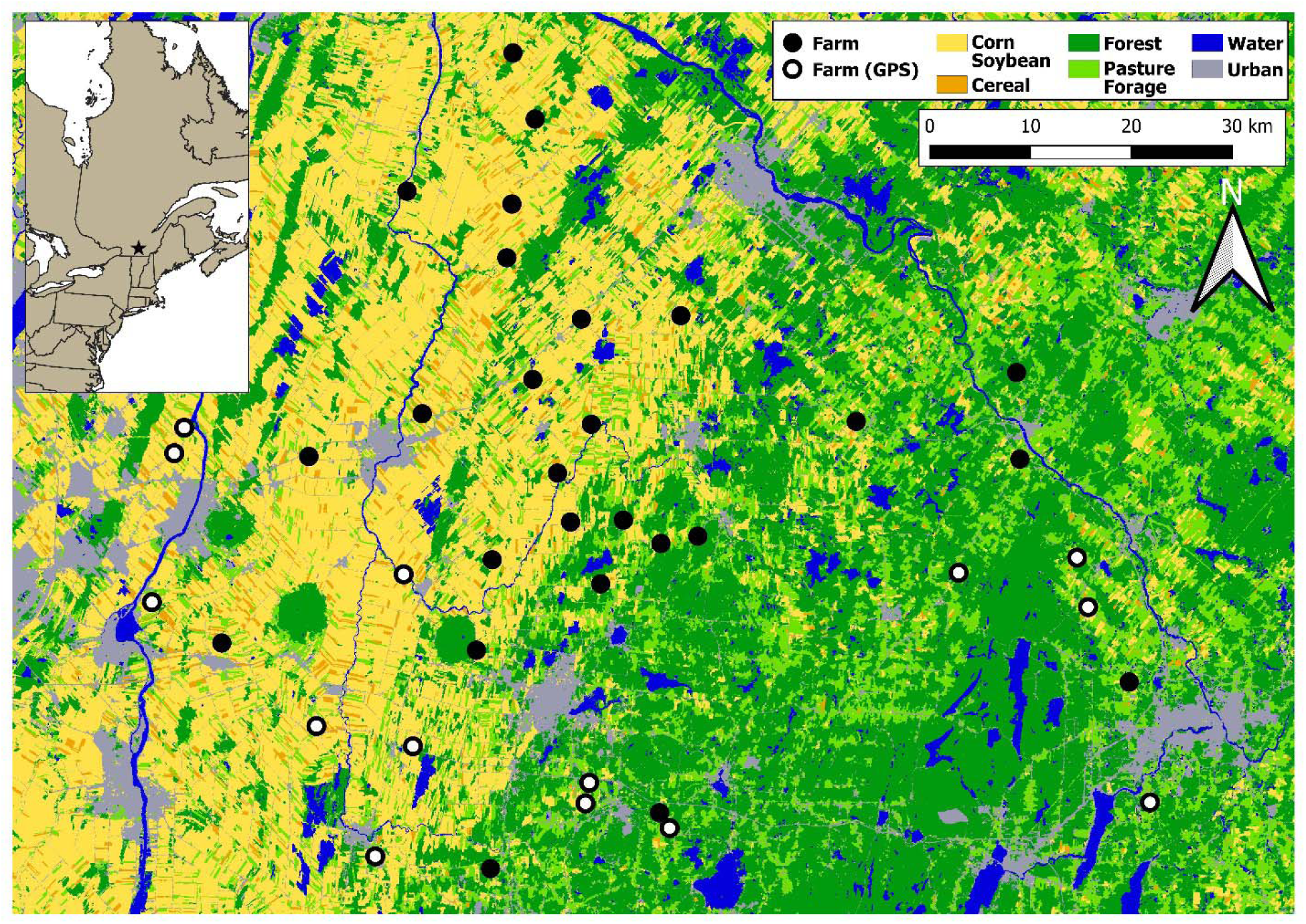
Map of study system. Each point represents the location of a farm, filled white points are farms where GPS were applied to 43 females. Underlying image represents the agricultural gradient derived from the Annual crop inventory (AAFC 2013). Light yellow represents the more agro-intensive areas while light green represents less agro-intensive areas.

## Methods

### Study sites and nest box visits

Data were collected between 2011 and 2016 within a nest box system placed across 40 farms throughout a gradient of agricultural intensification in southern Québec, Canada (Fig. 1, but see Ghilain and Bélisle 2008 for details). The western portion of the study area is composed primarily of large-scale row cropping, resulting in expanses denuded of forest cover and dominated by corn and soybean monocultures, and to a lesser extent, other cereals (principally wheat, *Triticum spp.*) (Bélanger and Grenier 2002, Ruiz and Domon 2009). This focus on monocultures has led to the near ubiquitous use of fertilizers, herbicides, insecticides and fungicides, as well as increased mechanization and removal of both surface waters and wetlands in this region (Jobin et al. 1996). These agro-intensive landscapes transition to ones composed primarily of pastures and forage crops (e.g. hay, alfalfa (*Medicago sativa*) and clover (*Trifolium spp.*)) in the eastern parts of the study system and are interspersed within large areas of forest cover and a systematic drop in pesticide detections (Poisson et al. 2021).

Ten nest boxes were placed on the field margins of each farm (total N=400) and have been monitored every two days throughout each Tree Swallow breeding season since 2006 (Porlier et al. 2009). The number of eggs and nestlings were counted during each nest box visit, and thus the age and phenology of each brood is known within two days of accuracy. We captured > 99% of adult females (during incubation) and >80% of males (during the brooding period) of each breeding attempt. Adults were banded with an aluminum US Geological Survey band containing a unique identification code and a series of morpho-metrics and samples were collected (Ghilain and Bélisle 2008, Lessard et al. 2014).

### Food provisioning

Food provisioning was characterized via two rates defining the amount and frequency of food being delivered to growing broods: the total biomass of delivered food boluses per hour (bolus biomass; mg/hr) and the number of food boluses delivered per hour (bolus delivery). These rates were estimated during food-provisioning sessions conducted between 2011 and 2016. Sessions consisted of two consecutive 30-min blocks in which collars were applied to each nestling within broods aged 6, 8, 10, and 12 days old (Bellavance et al. 2018). Collars were composed of a rubber band slid through a 5-mm long feeding tube and adjusted like a bolo-tie to capture food boluses in the crop yet loose enough to not restrict respiration. This method has often been used to document the diet of Tree Swallows, and has not been found to impact nestling survival (McCarty and Winkler 1999, Johnson and Lombardo 2000, Smits et al. 2005, Bellavance et al. 2018). At the end of each block, boluses were carefully removed via forceps and the nest was checked for discarded boluses. Boluses were then placed in sterile 50-ml conical tubes and immediately placed in a portable freezer (< 12 hours) until being stored at -80°C. At the end of each session, collars were removed and the starting time, the number of days past 1 January (Ordinal date) and the presence of precipitation was recorded. The maximum temperature during sessions was also noted at the end of each breeding season via iButtons (model DS1922L; Embedded Data Systems, Lawrenceburg, Kentucky, USA) attached to the underside of a single nest box on each farm that recorded hourly temperatures.

In a tandem study occurring between 2013 and 2016, the food boluses collected during food provisioning sessions were analyzed for the presence of the principal active agents of many pesticides (Haroune et al. 2015, Poisson et al. 2021). The methods of this study required recording the wet mass of food boluses (± 1 mg). We subsequently used this measure to estimate the total biomass of food boluses delivered during the one-hour food provisioning sessions.

### Foraging distance

The total distance an individual travels does not necessarily provide information on how far it ventured from its nest box. For central-place foragers, this distance will be related to the rate or effort to return to their nest box, and thus to the rate of food provisioning. Therefore, considering both the distance an individual travels and how far it ventured from its nest box will be more indicative of the possible foraging costs incurred than either value by itself. Consequently, we used GPS locations of nestling rearing females (nestling age 4-12 days) to estimate their total hourly distance traveled (DT) and mean hourly distance from their nest box (DNB). These metrics were assumed to be indicative of the amount of travel undergone by each individual and therefore to reflect the amount of energy expended while food provisioning. We used hourly travel distances to have a comparable time scale to the hourly rates estimated by the food-provisioning sessions. Specifically, we fitted PinPoint-10 GPS tags (Lotek Wireless Inc, Newmarket, Ontario, Canada) to 43 nestling rearing females across 14 farms in 2019 (Fig. 1, white points). Chosen farms represented landscapes dominated by either forage and pasture fields or the principal row-crop monocultures of our study system (i.e., corn, soybean and other cereals), with minimal tree cover (Appendix S2: Table S1). Tags weighed between 0.9 g and 1.2 g, including the harness (Appendix S2: Fig. S1), and were fitted over the synsacrum using leg loops (Rappole and Tipton 1991). We preprogrammed tags so GPS fixes (“SWIFT” fixes, nominal horizontal accuracy ± 10 m) were taken every 15 min between 5:00 and 20:00 local time (N = 60 locations) the day after deployment, allowing for a habituation period. This schedule was the most intensive that we could use to cover the majority of the daily food provisioning period given the battery-life (McCarty 2002, Rose 2009). Because of the amount of time elapsed between each GPS location, we did not intend for these locations to depict the actual path that individuals traveled throughout the day, and instead assumed that they provide enough information to derive a proxy of the total distance an individual may have traveled. We assessed the validity of this assumption by comparing GPS locations collected every 15 minutes to those collected every 5 minutes for two females (Appendix S2 Fig S2 and Fig S3). We observed that the total travel distance logically increases, and yet no general pattern was missed in terms of the distance traveled or relative frequency distribution of distances between GPS locations and an individual’s nest box. However, recording locations at this frequency severely impacted the longevity of the tag, resulting in the acquisition of GPS locations from only half the day when compared to sampling every 15 minutes. Given that sampling every 15 minutes did not result in the loss of major foraging forays, we are confident that this degree of sampling is the most ethically possible given technological constraints. The day after data collection, individuals were caught, and their tag removed. Once data were extracted, movements of each individual were derived by connecting GPS locations, in chronological order, and then calculating the distance along each segment of the resulting path using both the sp (Bivand et al. 2013) and rgeos (Bivand and Rundel 2019) packages in the R interface (R Core Team 2019). We estimated DT by summating the length of each segment between each point during each one-hour sampling period. We estimated DNB by first calculating the distance between the nest and each point, and then calculated the mean distance of these points within each one-hour block (Appendix S2: Table S2 and Fig. S4).

### Landscape characterization

Landscape characterization started with delineating habitats, including agricultural parcels, within 500 m of each nest box using orthophotos (scale 1:40 000) in QGIS (QGIS 2020) at the end of each breeding season between 2011 to 2019. This spatial scale hosts close to 80% of nestling rearing females’ daily time budget. We then characterized each habitat parcel in situ, defining agricultural habitats by the cultures grown. We then reclassified habitats into one of four higher order categories represented by forest, corn and soybean, forage fields (including pastures, hay fields, alfalfa, and clover), and cereals (principally wheat). Higher order categories represent the hypothesized habitats Tree Swallows use as indicators during nest-site selection, and the principal habitats composing landscapes throughout this system (Rendell and Robertson 1990, Courtois 2020, Winkler et al. 2020).

We calculated the mean percent cover of each higher order category across all nest boxes on each farm for each breeding season, resulting in 240 separate landscapes (i.e., 40 farms x 6 years). To obtain an integrative measure of the percent cover of all higher order habitats, defined as the landscape context, we used a robust principal components analysis (PCA) for compositional data (Filzmoser et al. 2009) to assign “site scores” to each of the farms during each year. Site scores were defined by the values along the first two components of the resulting compositional PCA as fitted using the robCompositions package in R (Templ et al. 2011). The first component (Comp.1) explained 79.8% of the variance in landscape context and was correlated positively with corn and soybean cultures and negatively with both forest cover and forage fields (Fig. 2A). The second component (Comp.2) explained 16.3% of the variance and was positively correlated with forest cover, negatively with both forage and cereal covers. Site scores were assigned to each food-provisioning session throughout the study.

**Figure 2:**
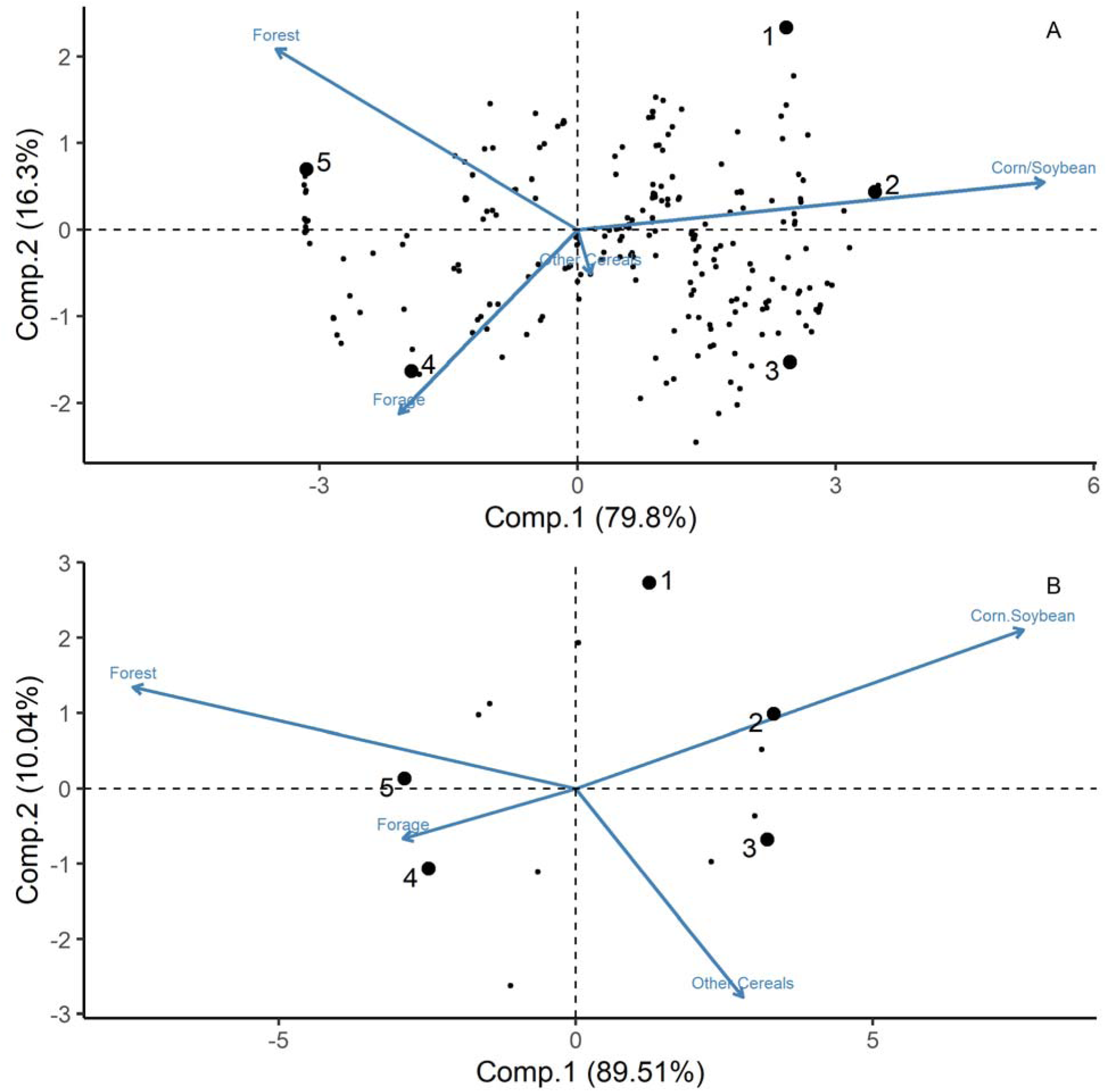
Results of the robust PCA for compositional data including the site scores of the five landscape context scenarios for the food provisioning sessions (A) and foraging distance studies (B). Points in the background represent the site scores assigned to each food provisioning session. The arrows lengths are eigenvectors representing the direction and magnitude of correlation between habitats. The five landscape scenarios are: ones dominated by corn or soybean with 8.8% forest cover (Corn/soybean with forest, 1), ones denuded of forest cover and dominated by only corn or soybean (Corn/Soybean,2), ones denuded of forest cover and dominated by a mixture of corn, soybean and cereals (mixed row crop, 3), ones minimizing forest cover (9.0%) and dominated by forage crops (Forage, 4), and agricultural landscapes with 69.0% forest cover (Agro-forested, 5)

The study of foraging distance occurred separately and on a subset of farms than those used to estimate food provisioning (Fig. 1). We thus repeated the landscape characterization used in the food-provisioning study, yet for the subset of farms hosting the foraging distance study. The resulting PCA was similar to that of the food provisioning sessions. The first component explained 89.5% of the variance in landscape context and correlated positively with both corn and soybean cultures, and negatively with both forest and forage cover (Fig. 2B). The second component explained 10.8% of the variance and correlated positively with forest cover as well as with both corn and soybean cultures, and negatively with forage and cereal cultures.

For ease of interpretation, all statistical predictions and interpretations were made in reference to five landscape context scenarios. These scenarios represent the predominant types of landscape contexts experienced across the agricultural gradient and are defined by the site scores of five farms (1-5) identified in Fig. 2A and B. The scenarios for the food-provisioning sessions are: (1) ones dominated by corn or soybean with 8.8% forest cover (corn/soybean with forest); (2) ones denuded of forest cover and dominated by only corn or soybean (corn/soybean); (3) ones denuded of forest cover and dominated by a mixture of corn, soybean and cereals (mixed row crops); (4) ones with 9.0% forest cover and dominated by forage crops (forage); and (5) landscapes with 69.0% forest cover and few cultivated parcels typically devoted to forage crops (agro-forested). Although site score values differ between the PCA derived for the food-provisioning and foraging distance studies, they are qualitatively similar and the above landscape context scenarios apply to both.

The distance of GPS locations from their respective nest boxes, greater than 0.5 km from nest boxes (∼20% of locations), was positively related to the percent cover of water bodies (e.g. rivers, wetlands and ponds) around a point (Appendix S2: Fig. S5 and Fig. S6), supporting the contention that these features are important (Elgin et al. 2020, Berzins et al. 2020). These habitats are however rare within our system and were considered as potential confounding factors of the observed foraging responses. Moreover, we hypothesized that the influence of water features located far from a nest may depend on their local availability (i.e., close to the nest). We therefore undertook a modeling endeavor aimed at identifying the spatial extent and the type of water features most influential to the rate of food provisioning (Appendix S3: Fig. S1).

Due to several model convergence problems, we found numerical stability was greatest when treating water features as open bodies of water (hitherto referred to as surface water). A local maximum of AICc weight (*w*) was observed when including the percent cover of surface water calculated within 1 km (local percent cover) and 10 km (regional percent cover) from the nest including their interaction. These model terms were included in all models for all foraging proxy. Surface water data were derived from vector layers acquired from the Canadian National Hydro Network data sets (NHN 2020) and Ducks unlimited (Ducks Unlimited, 2020). All calculations of percent cover were made using the sf (Pebesma 2018) R package.

### Local prey availability

Local prey availability was represented by the dry biomass of Diptera found within insect samples from two separate insect traps placed on each of the 40 farms between 2006 and 2016. We focused on dry biomass of Diptera (using the number of Diptera provides qualitatively similar results, analyses not shown, see also Rioux Paquette et al. 2013, Garrett et al 2021) because they represent a majority of the insects found in the diets of nestling Tree Swallows (McCarty and Winkler 1999, Johnson and Lombardo 2000, Mengelkoch et al. 2004, Twining et al. 2018). Within our study system, 74% of the biomass and 77% of the insects found in food boluses delivered to nestlings are Diptera (Rioux Paquette et al. 2013, Bellavance et al. 2018, Garrett et al. 2021a). The focus of this work being on the availability of food during the nestling period, we processed insect samples collected between 1 June and 15 July, representing the nestling period of over 96% of first breeding attempts. Diptera within each insect sample were removed, counted, and then placed in an oven at 50°C for over 24 hours to ensure no further changes in mass. Sample dry mass was then recorded (± 0.0001 g) without delay.

To obtain prey estimates more reflective of the farm vs. local scale of each trap, and avoid punctuated events such as insect swarms, we chose to use daily model predictions of Diptera biomass from generalized additive models (GAMs) in which raw values were regressed against the ordinal date of sample collection for each farm and year using a Gamma distribution and log link function. GAMs were fitted as a tensor product smoother using the mgcv package in R (Wood 2017) with an identical degree of smoothness (k=10). Model predictions are the accumulated biomass of Diptera over a roughly 48-hour period occurring over a 3-day window (i.e., from the time the trap was reset on day one, the entirety of day 2, and until the content of the trap was collected on day 3). As we wanted estimates of Diptera biomass on the day of the food provisioning session, prey availability is represented by the predicted Diptera biomass on the day following the food provisioning session.

### Statistical methods

#### Food provisioning

Bolus biomass was a continuous right-skewed (conditional) response containing zeros and was consequently modeled with generalized linear mixed effect models (GLMM) using a tweedie distribution and a log link function (Lo and Andrews 2015). We modeled bolus delivery with GLMM using a negative binomial distribution. In order to account for differences in sampling effort (both 30-min blocks were completed in 86% of sessions), we included whether sessions lasted half or an entire hour as an offset (i.e., log(0.5) or log(1), respectively) in both sets of analyses. We further included year, farm, and brood IDs as random factors in order to account for the data’s hierarchical structure (i.e., broods nested within farms nested within years).

We took an information theoretic and multimodel approach to evaluate links between landscape context and rates of food provisioning by comparing a set of 12 models representing biologically relevant alternative hypotheses (Table 1 and Appendix S2: Table S3). Except for a Null model containing only random effects, we started with a Base model including potential confounding variables. This model contained two weather variables. First, the mean maximum temperature, as insects require a threshold temperature before being active (Taylor 1963, Grüebler et al. 2008), and the upper limit of foraging swallow behavior is potentially thermally constrained (Tapper et al. 2020). Second, the presence of precipitation, as we expected precipitation to act as a hindrance to foraging (Cox et al. 2019), and because food provisioning sessions did not last long enough to estimate amount of precipitation. The base model also included the age of the female (second year (SY) or after second year (ASY); Hussell 1983), time at the start of the provisioning session as a second-order polynomial, the percent cover of surface water within 1 km and 10 km and their interaction (Elgin et al. 2020), and a proxy of the brood’s food demand (Geary et al. 2020). This brood demand proxy was represented by the multiplication of the brood size with the brood age. We did not expect foraging rates to respond linearly with this proxy, as the functional response likely reaches a plateau whereby food provisioning parents can no longer increase their efforts. We therefore treated this variable as a third-order polynomial. All model terms within Base were included in all subsequent models of the candidate set. Our principal model of interest was one in which sites scores and their interaction were added to assess landscape context effects (Base + Land). Because potential negative effects of foraging within agro-intensive landscapes may be alleviated with increasing prey availability closer to nests, we also considered models with only local prey availability (Base + Food), one with both prey availability and site scores (Base + Food + Land), and one also including the interaction between prey availability and Comp.1 (Base + Food*Land). We further hypothesized provisioning rates may be influenced by brood demand and the availability of prime foraging habitat and included four models with such relationships. These included one in which the effect of land cover and local prey availability varied with the demand proxy (Base + Food + Land * Demand and Base + Land + Food * Demand) or the local (1 km) percent cover of surface water (Base + Land + Food * LSW and Base + Food + Land * LSW).

**Table 1:** Mean ± SD of covariates used to model response variables. Covariates for the hourly number of boluses and total bolus biomass delivered to nestlings were representative of the hourly food provisioning sessions conducted between 2011 and 2016. Covariates of foraging distances were calculated for the hourly interval’s representative of the hourly GPS locations. Dashes imply the covariate was not used in the modeling of the response variable. Covariate groups identify which covariates were present within models found in Table 1.

| Covariate | Acronym | Unit | Covariate group | Biomass | Number | Foraging distance |
| --- | --- | --- | --- | --- | --- | --- |
| Starting time | ST | Time | Base | 262.31 ± 150.16 | 251.57 ± 153.55 | - |
| Maximum temperature | TMAX | °C | Base/Food | 26.20 ± 4.08 | 25.54 ± 3.94 | 20.81 ± 5.11 |
| Brood demand proxy | DM | Brood size *<br>Brood age | Base/Demand | 40.50 ± 13.89 | 39.14 ± 12.8 | 28.9 ± 10.59 |
| Local surface water (1 km) | LSW | % | Base/ LSW | 0.80 ± 1.43 | 0.87 ± 1.58 | 1.52 ± 1.73 |
| Regional surface water (10 km) | RSW | % | Base | 1.58 ± 1.46 | 1.56 ± 1.42 | 2.67 ± 1.93 |
| Prey availability (Diptera biomass) | PA | g | Food | 0.02 ± 0.01 | 0.02 ± 0.01 | - |
| Site score Comp.1 | Comp.1 | - | Land | 0.34 ± 1.73 | 0.42 ± 1.72 | 0.09 ± 2.42 |
| Site score Comp.2 | Comp.2 | - | Land | -0.04 ± 0.88 | -0.01 ± 0.86 | -0.25 ± 1.24 |
| Ordinal day | OD | days | Base | - | - | 170.64 ± 7.06 |

#### Foraging distance

Two response variables characterized the foraging distances of females, the total hourly distance traveled (DT) and the mean hourly distance an individual was from its nest box (DNB). These distances were highly correlated (*r*=0.8), yet considering both was ultimately more informative of foraging behavior. These response variables were modeled using linear mixed effects models (LMM). Both response variables were log-transformed to ensure normality of residuals. We included the individual and farm IDs as random effects to account for the hierarchical structure of the dataset (i.e., locations nested within individuals nested within farms).

We applied an information theoretic and multimodel approach to a set of six models to evaluate hypotheses about foraging distances (Table 1 and Appendix S2: Table S3). We started with a Null model only including random effects and a Base model including potential confounding variables influential to foraging distances (McCarty 2002, Dreelin et al. 2018, Elgin et al. 2020). Confounding factors included the brood demand proxy as a third-order polynomial, as well as the percent cover of surface water within 1 km and 10 km and their interaction. We further included proxies of local prey availability, including the ordinal date and the maximum observed temperature during each one-hour sampling period as a second-order polynomial (Rioux Paquette et al. 2013, Bellavance et al. 2018, Garrett et al. 2021a). Though the principal model of interests was one including site scores and their interaction (Base + Land), we compared another three models hypothesizing that the effects of foraging within agro-intensive landscapes may either be alleviated or exacerbated by confounding variables. We thus considered models also including an interaction term between Comp.1 and each potential confounder, namely brood demand (Base + Land*Demand), local (1 km) percent cover of surface water (Base + Land*LSW), and local prey availability (Base + Land*Food).

In all analyses, the effects of key individual variables, including interactions, were estimated via multi-model inference whereby predictions were calculated by model-averaging with shrinkage and shown with their 95% unconditional confidence intervals (Burnham and Anderson 2002). All quantitative covariates were z-transformed and all analyses were performed in R version 3.6.2 (R Core Team 2019) using the glmmTMB (Brooks et al. 2017) and AICcmodavg (Mazerolle 2020) packages. Model validation was performed following Zuur et al. (2009) using the DHARMa package (Hartig 2020).

## Results

### Food provisioning

#### Bolus biomass

We measured the total bolus biomass delivered during 741 food provisioning sessions from 420 broods, 92% of which consisted of both 30-min blocks. Overall mean bolus biomass was 284.5 mg per hour (SD = 344.9), ranging from 208.2 mg to 319.2 mg across years (Appendix S2: Fig. S7). The Null and Base models were not supported by the data, contrary to models including information on landscape context (Appendix S2: Table S3). Compared to forage landscapes (#4 in Fig. 3B and Appendix S2: Fig. S8B), bolus biomass was 54.0% lower within agro-forested landscapes (#5), 42.8% lower in mixed row-crop landscapes (#3), and 32.8% lower within corn/soybean landscapes (#2). Unlike for bolus delivery rate (see below), we did not find that the effect of corn/soybean cover was dependent upon forest cover (#1). We found that bolus biomass was unrelated to local prey availability as measured (Table 2 and Appendix S2: Fig. S9 and S10). We further found bolus biomass increased almost linearly with brood demand (Appendix S2: Fig. S10 and S11), and was negatively correlated with the local percent cover of surface water and was greater for ASY females (Appendix S2: Fig. S12). Lastly, bolus biomass increased with both the start time and the maximum temperature of the food provisioning session, until plateauing at 14:12 and 25°C, respectively (Table 2 and Appendix S2: Fig. S10).

**Figure 3:**
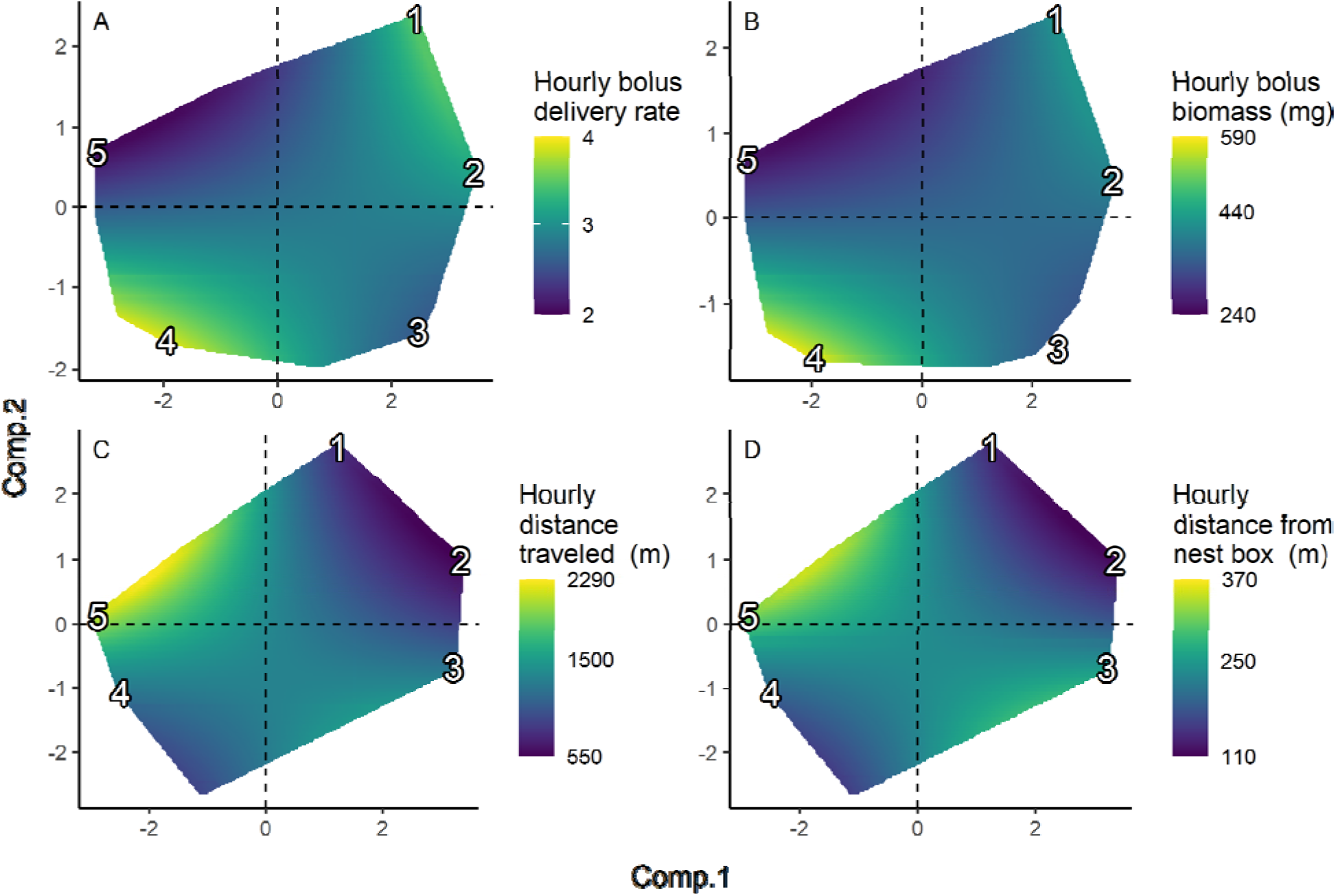
Response surfaces of the principal response variables against the first two components deriving the landscape context. Response surface are the two-dimensional unconditional predictions of each response variable over the observed range of values along the first and second component values of the PCA defining landscape context found in Fig 3. Deeper blue represents lower predicted values while yellow represents greater values. Predicted hourly bolus delivery rate (A), hourly bolus biomass (B) delivered to nestlings. Predicted hourly distance traveled (C) and hourly distance from nest box (D). Numbers in each panel are landscape context scenarios and represent the predominant types of landscape contexts experienced across the agricultural gradient and correspond to the points in numbers in Fig. 2.

**Table 2:** Standardized coefficient estimates and their 95% confidence intervals included in the top model of each response variable. See Table 1 for definitions of covariates and Appendix S2: Table S3 for outcome of model selection. Coefficients in bold indicate confidence intervals not overlapping zero. Numbers next to acronym refer to the order of the polynomial term.

| Covariate | Biomass | Delivery | Travel | Distance |
| --- | --- | --- | --- | --- |
| Comp.1 | **0.04**<br>(0.00, 0.08) | 0.02<br>(-0.04, 0.07) | **-0.11**<br>(-0.16, -0.05) | -0.08<br>(-0.16, 0.01) |
| Comp.1:Comp.2 | **0.08**<br>(0.03, 0.13) | **0.09**<br>(0.02, 0.16) | **-0.17**<br>(-0.25, -0.09) | **-0.15**<br>(-0.25, -0.05) |
| Comp.2 | **-0.11**<br>(-0.20, -0.02) | **-0.15**<br>(-0.27, -0.02) | 0.05<br>(-0.06, 0.16) | 0.03<br>(-0.15, 0.20) |
| Female Age (SY) | 0.09<br>(-0.09, 0.26) | 0.13<br>(-0.13, 0.40) | - | - |
| DM1 | **5.09**<br>(2.62, 7.55) | **9.78**<br>(7.09, 12.47) | **3.12**<br>(0.41, 5.82) | 3.21<br>(-0.92, 7.33) |
| DM2 | **-6.09**<br>(-8.58, -3.60) | **-3.34**<br>(-6.11, -0.57) | **-3.02**<br>(-5.81, -0.23) | **-4.76**<br>(-8.81, -0.71) |
| DM3 | **3.72**<br>(1.15, 6.29) | 1.07<br>(-1.64, 3.78) | -1.41<br>(-1.45, 4.26) | 1.45<br>(-3.02, 5.91) |
| DM1:Comp.1 | 0.47<br>(-0.90, 1.85) | - | - | - |
| DM2:Comp.1 | **1.65**<br>(0.17, 3.13) | - | - | - |
| DM3:Comp.1 | -0.65<br>(-2.29, 0.98) | - | - | - |
| OD1 | - | - | **10.80**<br>(7.32, 14.28) | **12.50**<br>(7.03, 17.97) |
| OD2 | - | - | **-5.62**<br>(-8.57, -2.67) | **-6.28**<br>(-10.90, -1.66) |
| ST1 | **2.93**<br>(0.53, 5.33) | 2.53<br>(-0.19, 5.26) | - | - |
| ST2 | **-2.49**<br>(-4.90, -0.09) | **-3.02**<br>(-5.61, -0.43) | - | - |
| TMAX1 | 0.87<br>(-1.62, 3.36) | 0.13<br>(-2.58, 2.83) | **8.79**<br>(6.34, 11.24) | **8.24**<br>(5.53, 10.96) |
| TMAX2 | **-5.77**<br>(-8.21, -3.32) | **-3.99**<br>(-6.75, -1.23) | **-5.35**<br>(-7.73, -2.96) | **-4.61**<br>(-7.23, -1.99) |
| Rain (Yes) | -0.17<br>(-0.44, 0.10) | -0.18<br>(-0.52, 0.16) | - | - |
| LSW | **-0.10**<br>(-0.17, -0.03) | **-0.12**<br>(-0.22, -0.02) | 0.17<br>(-0.02, 0.37) | 0.06<br>(-0.22, 0.35) |
| LSW:Comp.1 | - | - | **-0.09**<br>(-0.17, -0.02) | - |
| LSW:RSW | 0.03<br>(-0.07, 0.14) | 0.12<br>(-0.03, 0.28) | **-0.29**<br>(-0.52, -0.07) | -0.29<br>(-0.62, 0.05) |
| RSW | 0.06<br>(-0.02, 0.14) | 0.05<br>(-0.06, 0.16) | **-0.26**<br>(-0.44, -0.09) | **-0.31**<br>(-0.58, -0.04) |

#### Bolus delivery

We conducted 1,208 food provisioning sessions from 662 different broods. Overall mean bolus delivery was 2.00 boluses per hour (SD = 2.05), ranging from 1.69 to 2.25 across years (Appendix S2: Fig. S7). While the Null and Base models were not supported by the data, several alternative models received support (Appendix S2: Table S3). We observed substantial landscape effects (Table 2, Fig. 3A). Compared to forage landscapes (#4 in Fig. 3A), bolus delivery rate was 46.2% lower within agro-forested landscapes (#5), 34.5% lower in mixed row-crop landscapes (#3), and 19.3% lower within corn/soybean landscapes (#2). The effect of corn/soybean cover was however highly dependent upon forest cover as the presence of remnant woodlots in landscapes highly dominated by these crops (#1) was associated with a bolus delivery rate 11.2% lower than within forage landscapes (#4). We found that bolus delivery rate was not related to local food availability (Appendix S2: Fig. S9 and S13). We further found that bolus delivery rate was positively associated with brood demand (third-order) and both the start time and the maximum temperature of the food provisioning session, until plateauing at 14:59 and 26°C, respectively (Appendix S2: Fig. S11 and S13). It also negatively correlated with the local percent cover of surface water (Appendix S2: Fig. S14).

### Foraging distance

We successfully deployed and recovered 43 GPS tags. Of these tags, 40 successfully recorded all 60 attempted locations, resulting in 2,516 swift-fixes and 631 hourly estimates of foraging distances (Appendix S2: Table S2 and Fig. S4). The overall mean distance of each location from an individual’s nest box was 420 m (SD = 930) and 465 m (SD = 814) between each consecutive fix. Six females recorded locations over 5 km from their nest box with a maximum distance of 9.5 km.

#### Hourly distance traveled

Overall mean DT was 1,857 m (SD=2,365, Appendix S2: Fig. S7). A single most predictive and parsimonious model occurred when considering landscape context along with an interaction between Comp.1 and local percent cover of surface water (Table 2 and Appendix S2: Table S3). DT was longest within agro-forested landscapes (#5 in Fig. 3C) and was 75.6% further when compared to forage landscapes (#4). Compared to forage landscapes, distances were 2.8% shorter within mixed row-crop landscapes (#3), and yet 52.7% shorter within corn/soybean landscapes (#2). The effect of corn/soybean cover was however highly dependent upon forest cover, as the presence of remnant woodlots in landscapes highly dominated by these crops (#1) was associated with a DT 34.2% shorter than within forage landscapes (#4). DT increased with brood demand until plateauing at mean demand values (Appendix S2: Fig. S11 and S15). DT was further associated in a quadratic fashion with both the ordinal date and maximum temperature, peaking on ordinal date 183 (2 July) and at 26.1°C (Appendix S2: Fig. S15). Finally, DT was shorter with increasing regional cover of water (Table S2, Appendix S2: Fig. S16).

#### Hourly mean distance from nest box

Overall mean DNB was 421 m (SD=719, Appendix S2: Fig. S7). While no single model could be considered the most predictive or parsimonious, models taking landscape context into account were clearly better supported (Table 2 and Appendix S2: Table S3). DNB was furthest within agro-forested landscapes (#5 in Fig. 3D) and was 65.5% further when compared to forage landscapes (#4). Compared to forage landscapes, DNB was 34.3% further within mixed row-crop landscapes (#3), and yet 37.4% closer within corn/soybean landscapes (#2). The effect of corn/soybean cover was however highly dependent upon forest cover as the presence of remnant woodlots in landscapes highly dominated by these crops (#1) was associated with DNB being 24.4% closer than within forage landscapes (#4). DNB increased significantly until plateauing at mean values of brood demand (Appendix S2: Fig. S11 and Fig. S17). DNB was associated in a quadratic fashion with both the ordinal date and maximum temperature, peaking on ordinal date 184 (3 July) and at 26.1°C (Appendix S2: Fig. S17). Finally, DNB was shorter with increasing regional cover of water (Table S2, Appendix S2: Fig. S18).

## Discussion

This study aimed at evaluating a hypothesized mechanism that negatively correlated the fitness of an aerial insectivore with breeding in landscapes increasingly composed of more agriculturally intensive monocultures (e.g., corn and soybean). This mechanism hypothesized that foragers in agro-intensive landscapes expend more time or energy reaching food patches and traveling between them, with the additional cost of settling for patches of lower quality, the combination of which would result in lower food provisioning rates (Appendix S1: Fig. 1). In more agriculturally dominated landscapes, we found that female Tree Swallows provided food to their nestlings at lower rates within increasingly agro-intensive landscapes. The hourly rate of delivery of both bolus biomass and bolus number were indeed lowest within more agro-intensive landscapes and greatest within forage landscapes. Female Tree Swallows traveled most and furthest from their nest within forest-dominated landscapes, and unexpectedly less and closer to their nest in agro-intensive compared to forage landscapes.

The relationship between the rate at which boluses were delivered and landscape context resembled that at which prey biomass was provided, suggesting that the biomass of individual boluses was similar throughout the agricultural intensification gradient studied (McCarty 2002). This outcome was corroborated by a supplemental analysis (Appendix S4: Table S1 and S2). Furthermore, coupled with the result that females traveled furthest within predominantly forested landscapes and to a lesser extent in forage landscapes, the relatively low variance in the biomass of individual boluses suggests that forage landscapes result in marginally greater travel times.

Yet the increased frequency of food bolus delivery implies that foraging in forage landscapes likely lead to more rapid loading curves, and greater gain rates, than that of agro-intensive landscapes (Fig. 4). Such results stand, even if we assume Tree Swallows from this system do not experience a decelerating gain curve while exploiting food patches, and instead “sift” through the air column until reaching an individual-dependent fixed loading capacity (Appendix S2. Fig S2).

**Figure 4:**
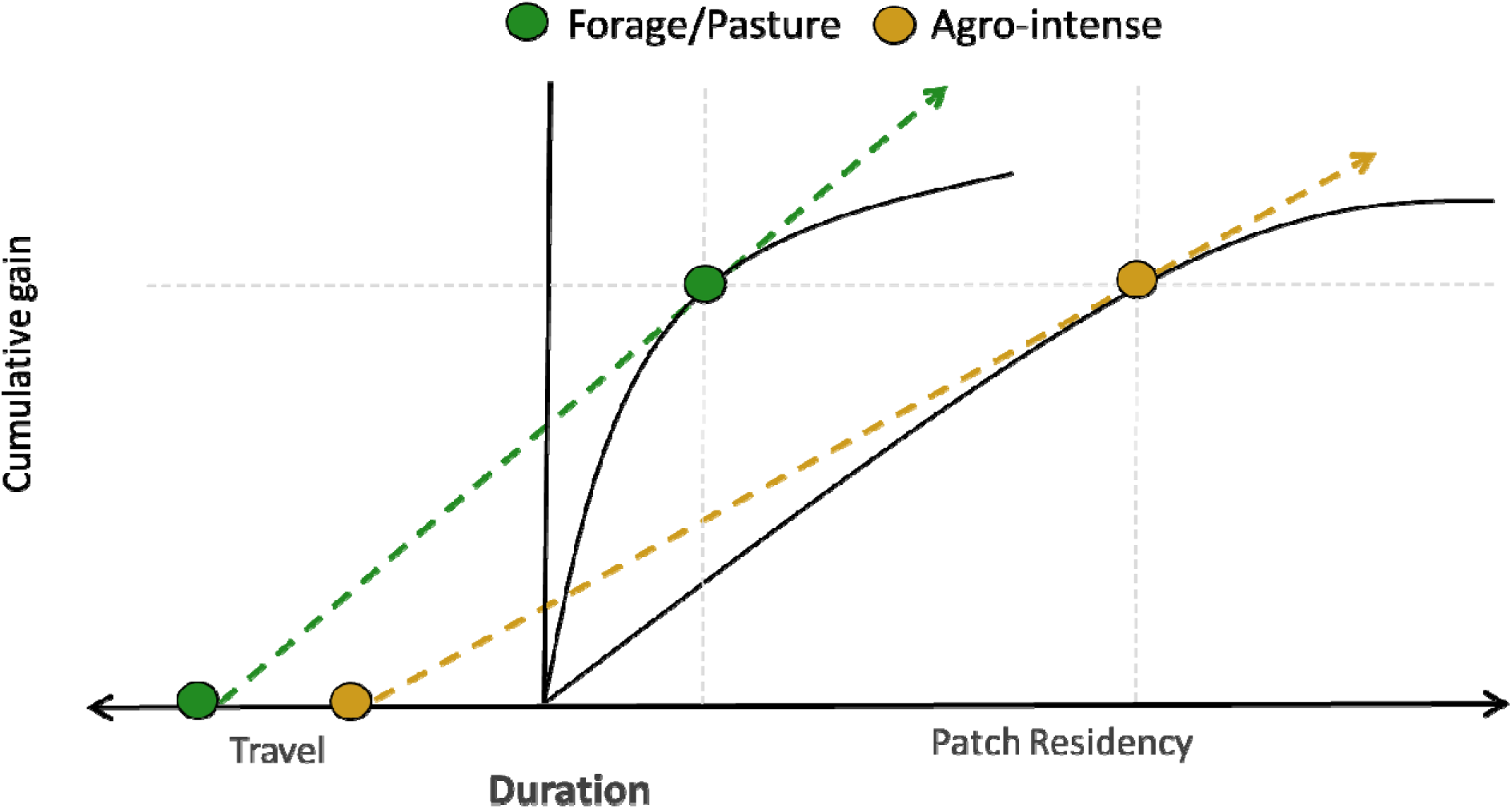
Reasoning underlying our hypothesis that food patches within forage landscapes allow for a more rapid accumulation of food resources than those found in more agro-intensive landscapes. Assuming that food provisioning Tree Swallow females foraged according to th marginal value theorem, and given we found (1) that the individual bolus biomass was similar across the agricultural intensification gradient, (2) that travel distances were greater in forage than in agro-intense landscapes, then only a lower gain curve in agro-intense landscapes is compatible with our findings that hourly bolus biomass and hourly number of boluses delivered were higher in forage than in agro-intense landscapes. Gain rate is graphically represented by the slope of the dashed line connecting mean travel time to the tangent of the gain curve.

It should be stated that our results were focused on correlations between landscape composition and food provisioning rates and not on how these rates are correlated to measures of avian fitness. Therefore, conclusions from these results should recognize the need to better establish causal pathways between provisioning rates and both nestling and adult body condition, as well as nestling survival. However, these factors are intimately linked, and despite our work being correlative, they are ultimately proxies of some underlying mechanisms warranting future work. For example, lower food provisioning rates within more agro-intensive landscapes may need to be compensated by parents elongating the entire foraging period of a given day, foraging more intensely when conditions allow, forgoing recuperation or foraging for themselves (Martins and Wright 1993, Markman et al. 2002, Ydenberg 2007, Schifferli et al. 2014). In all cases, increases in foraging effort may come at a cost to themselves through reduced body condition and thus potentially impactful to their future reproductive success (Stearns 1992, Reznick et al. 2000, Harrison et al. 2011, Lagrange 2015), as increases in effort may result in foraging parents being in too poor of body condition for molting prior to migration (Jenni-Eiermann and Jenni 1996, Rubolini et al. 2002). Furthermore, if parents cannot compensate for lowered food provisioning rates, then broods may incur the cost through reduced growth rates or elevated mortality (Naef-Daenzer et al. 1999, Tremblay et al. 2004). In fact, evidence suggest food provisioning Tree Swallows as well as their broods already incur these types of costs. For example, parents breeding in more agro-intensive landscapes spend less time in the nest (Lamoureux 2010, Stanton et al. 2016), express reduced body condition (Pigeon et al. 2013, Stanton et al. 2017), and both reduced fledging success and nestling growth (Houle et al. 2020, Garrett et al. 2021a). Some of these natal effects are of concern as they may be carried over into later life stages (Harrison et al. 2011).

One fundamental characteristic of our gradient of agricultural intensification, and as an extension, most agricultural landscapes, is a loss of forest cover (Fischer and Lindenmayer 2007). Within our study area, landscapes dominated by a mixture of forest cover and agricultural fields resulted in the lowest food provisioning rates and furthest foraging distances. The foraging habitat of Tree Swallows, as well as of most swallow species, is frequently recognized as being open terrestrial habitats or over large water surfaces (Evans et al. 2007, Boynton et al. 2020, Elgin et al. 2020). Thus, forest cover may reduce landscape functional connectivity for these species, at least with respect to foraging (Bélisle 2005, Olsson and Bolin 2014, Evens et al. 2018). In this way, as forest cover is reduced, the cover of foraging habitats, close to central places and conducive to Tree Swallows, increases. In fact, second to forage landscapes, we found provisioning rates were superior within agro-intensive landscapes with remnant forest cover. Residual woodlots within these agro-intensive landscapes add to the tree lines separating fields to increase the length of “forest” edges, where insect abundance tends to be greatest (Jokimäki et al. 1998, Evans et al. 2003, Boetzl et al. 2020). The fledging success of Tree Swallows from the system presented here is greater in agro-intensive landscapes containing residual woodlots than ones denuded of forest cover (Garrett et al. 2021a). It may therefore be hypothesized that the elevated insect abundance associated with these residual woodlots lead to greater provisioning rates and thus greater fledging success in this landscape context.

It should be noted that during the years covered by this study, areas intensively-managed for the growing of corn and soybean harbored elevated levels of pesticides in their surface waters, notably several herbicides (e.g. glyphosate, atrazine, S-metolachlor, imazethapyr) and neonicotinoid insecticides [e.g. clothianidin, thiacloprid, thiametoxam (Giroux 2019, Montiel-León et al. 2019)]. For instance, a previous study found that, on average, exposure to active pesticide agents within the food boluses was greatest within these landscapes (Poisson et al. 2021). The consumption or prolonged exposure to such chemicals can influence the neurological, endocrine and immunological systems of non-target animals, including vertebrates, via “skin” contact, breathing or consumption of food or water (Mayne et al. 2005, Mineau et al. 2013, Lopez-Antia et al. 2015, Pisa et al. 2015). Importantly, neurotoxic insecticides can induce anorexia, alter orientation and influence motor functions required for flight in birds (Eng et al. 2017, 2019). Foraging within agro-intensive landscapes increases exposure of foragers to these chemicals, likely reducing their motivation and ability to forage, and thus potentially resulting in decreased provisioning rates.

Finally, foraging is a metabolically expensive behavior, made worse when foraging bouts are during the nestling periods (Bryant and Tatner 2008, Yap et al. 2017). Therefore, incoming food need not only be frequent, but also nutritionally rich (Twining et al. 2018). The diversity and abundance of insects found within food boluses of this system suggests that the nutritive value of food varies across the agricultural gradient (Bellavance et al. 2018). Most notably, insects with aquatic life stages (principally Ephemeroptera) are more abundant within the food boluses retrieved from forage landscapes than from those dominated by more agro-intensive cultures.

These insects are greater in highly unsaturated omega-3 fatty acids (HUFA; (Twining et al. 2018). Nestlings with diets rich in HUFA can grow faster, gain a better body condition and have greater immunocompetence, leading to elevated brood survival and fledging success (Twining et al. 2018). Such effect can logically be extended to foraging adults. Subsequently, the energetic losses of foraging may be more easily replenished within forage landscapes.

To our surprise, we found that both measures of food provisioning were not related to our measure of local prey availability. Our measure of food availability was based on sampling over a two-day period prior to nest box visits, while our sampling of food provisioning occurs over a one-hour period. We believe that, despite food availability being linked to the fitness of Tree Swallows in this system (Garrett et al. 2021a), the differing temporal scales of sampling likely hinders our ability to confidently evaluate how local food availability is related to food provisioning. Instead, we suggest future work explicitly focus on sampling aerial insects during food provisioning sessions, as has been used in other similar studies (Grüebler et al. 2008, Stanton et al. 2016, Elgin et al. 2020).

Global climate change results in increased global temperatures and predicts increases in both the frequency and intensity of extreme weather events, such as prolonged periods of low or high temperatures or precipitation (Rahmstorf and Coumou 2011, Wuebbles et al. 2014). Such weather events may drastically alter the availability of insects as a food resource due to their thermally dependent nature (Grüebler et al. 2008, Shipley et al. 2020, Garrett et al. 2021a). Furthermore, recent evidence suggests Tree Swallows may already be foraging at some upper thermal threshold (Tapper et al. 2020). Our results, combined with other pieces of evidence, suggest food provisioning adults may also be foraging at some upper energetic limit inasmuch as proxies of body condition are lower where food provisioning rates are lower (Pigeon et al. 2013, Stanton et al. 2016, 2017). It is thus imperative that future work focuses not only on how agricultural practices or global climate change independently impact declining avian species, but also on how these factors may interact with one another. This importance is due to the ability of growing broods or breeding parents to manage extreme weather is potentially lower within more agro-intensive landscapes (Grue et al. 1997, Evans et al. 2003, Eng et al. 2019, Tapper et al. 2020, Garrett et al. 2021b). Finally, our results provide evidence that aerial insectivore declines may be related to agricultural intensification through a trophic pathway.

## Supporting information

AppendixS1

AppendixS2

AppendixS3

AppendixS4

## Acknowledgments

We are indebted to the farm owners who kindly accepted to partake in our long-term study since 2004. We greatly appreciate the time and effort of the reviewers. Sincere thanks to Josiane Côté Audet and the many graduate students, field and lab assistants who helped collect, enter, and proof data throughout the years, notably with respect to insect sample processing. This work was conducted under the approval of the animal care committee of the Université de Sherbrooke and was financially supported by Natural Sciences and Engineering Research Council of Canada discovery grants to FP, DG and MB, two team research grants from the Fonds de recherche du Québec—Nature et technologies to FP, DG and MB, by the Canada Research Chairs program to FP and MB, as well as by the Canadian Foundation for Innovation to FP, DG and MB, the Canadian Wildlife Service of Environment and Climate Change Canada and the Université de Sherbrooke.

