## AppendixS1 for "Negative effects of agricultural intensification on the food provisioning rate of a declining aerial insectivore"

**Appendix S1**

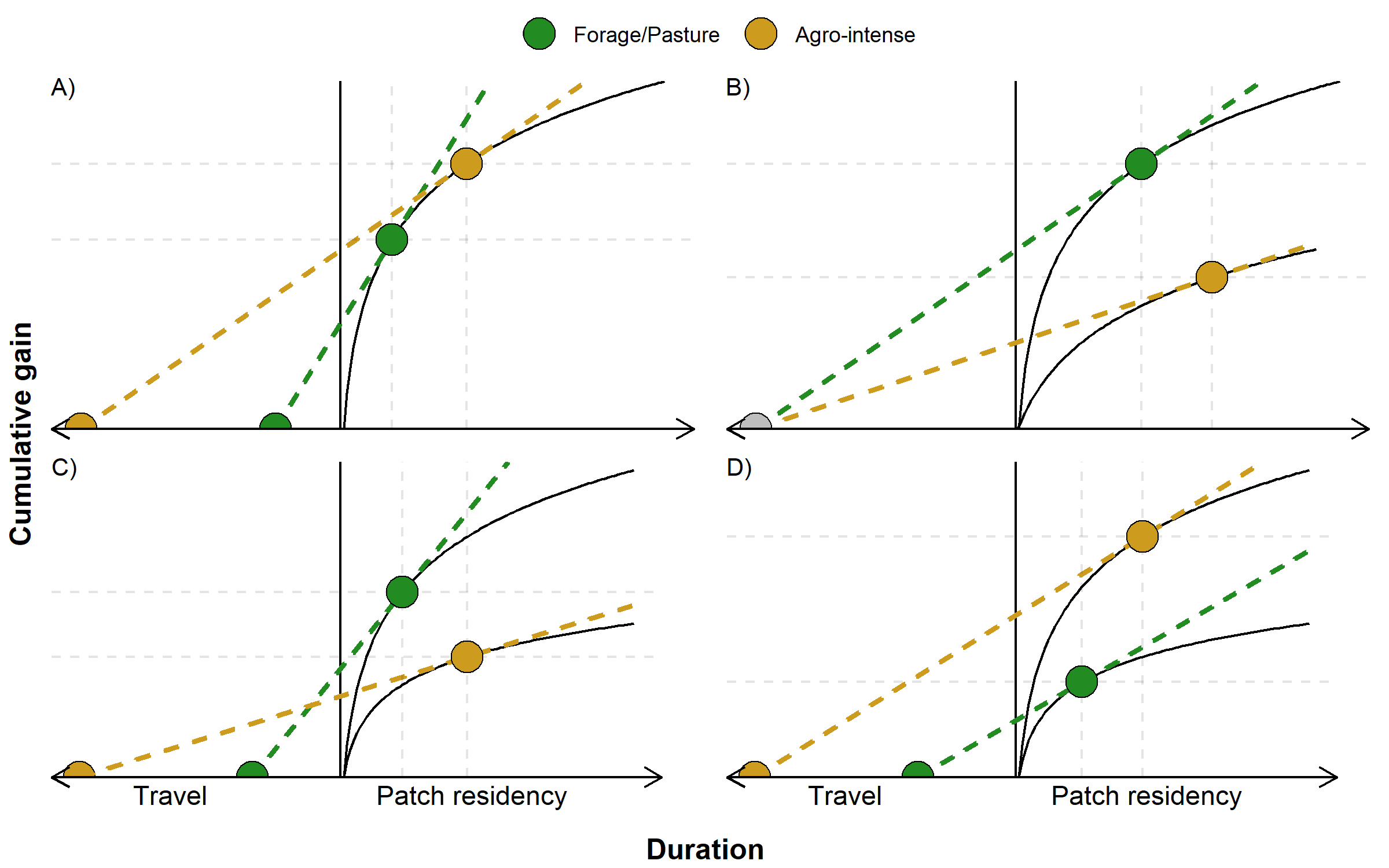

Figure S1: Optimal gain rates expected under different scenarios where food provisioning individuals forage according to the marginal value theorem and experience different travel times and gain curves that depend (or not) on landscape context. The x-axis is the total duration of a foraging bout composed of both the travel time to foraging patches and the time spent foraging within patches. Relationship between cumulative gain and patch residency time is represented by a decelerating gain curve. The food provisioning rate given a foraging bout is the ratio between the cumulative gain and the bout’s total duration. Optimal gain rate is graphically represented by the slope of the dashed line connecting travel time to the tangent of the gain curve. Assuming more agro-intensive landscapes result in more dispersed or poorer foraging patches, landscape mediated differences in food provisioning rate potentially occur through longer travel times (A and C) or lower instantaneous gain rates while exploiting patches (B and C). Lastly, shorter travel times can counterbalance slower gain rates within food patches, resulting in different food provisioning rates and yet similar gains per unit time between contrasting landscapes (D).

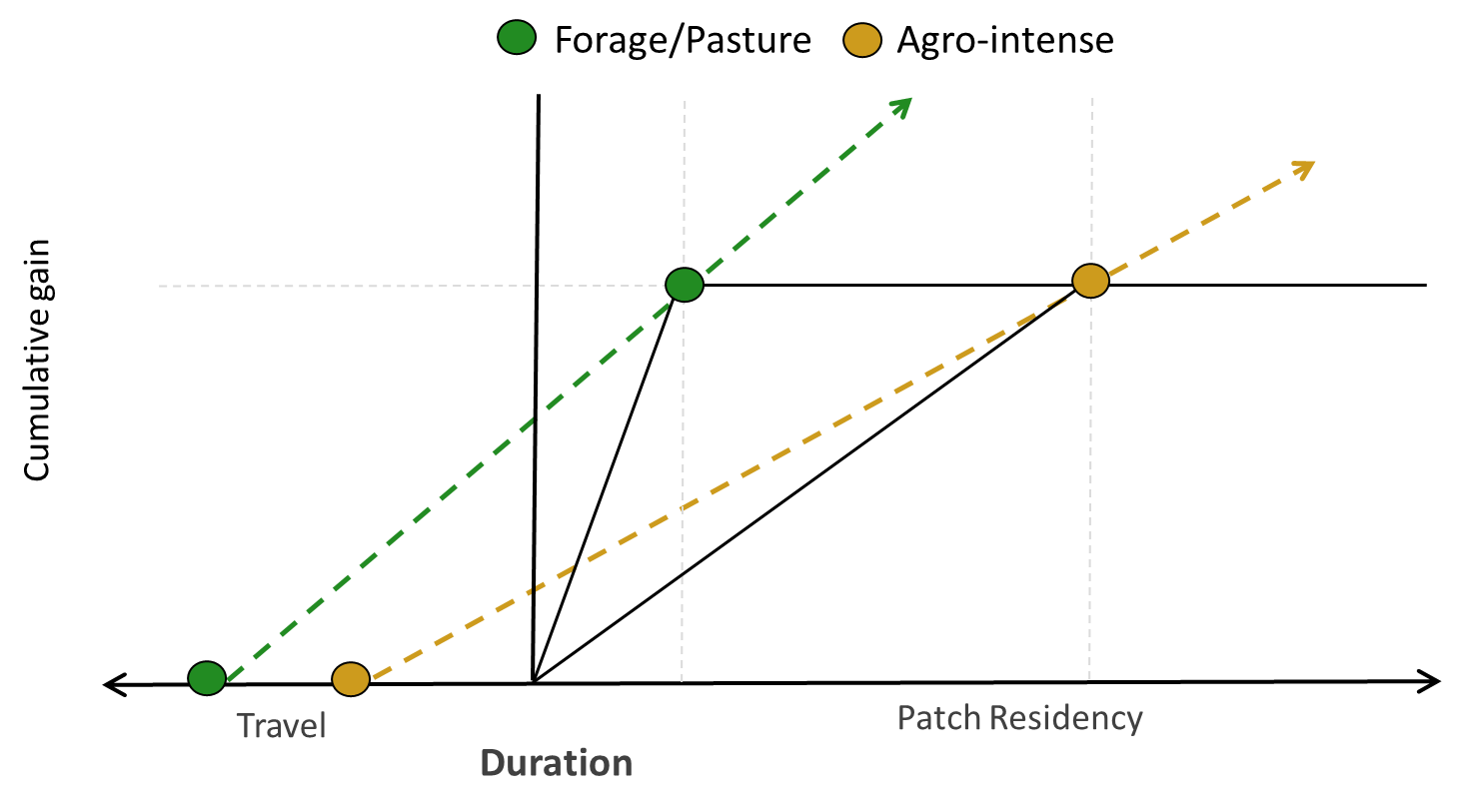

Figure S2: Our results indicate that individuals from this study system may not forage in discrete resource patches. Instead, individuals from this population may sift the air column and forage until reading an individual-dependent fixed loading capacity, before returning to their nest. Presented is she same information as in Figure 5 with the removal of the model constraint that the cumulative gain curve is a decelerating positive relationship, and further demonstrates our underlying hypothesis that foraging within more forage landscapes allow for a more rapid accumulation of food resources than in more agro-intensive landscapes. Assuming food provisioning Tree Swallow females foraged according to the marginal value theorem, and given we found (1) that the individual bolus biomass was similar across the agricultural intensification gradient, (2) that travel distances were greater in forage than in agro-intense landscapes, then only a lower gain rate in agro-intense landscapes is compatible with our findings that hourly bolus biomass and hourly number of boluses delivered were higher in forage than in agro-intense landscapes.
