## AppendixS2 for "Negative effects of agricultural intensification on the food provisioning rate of a declining aerial insectivore"

### Appendix S2

Table S1: Landscape metrics of the farms in which GPS tags were applied to 43 breeding female Tree Swallows. Values are the percent cover of each delineated habitat. Unless explicitly stated, percent cover was calculated within 500 m of nest boxes.

| Farm ID | Comp.1 | Comp.2 | Corn Soybean | Forage | Other Cereals | Forest | Water local (1 km) | Water regional (10 km) |
| --- | --- | --- | --- | --- | --- | --- | --- | --- |
| 01 | 1.24 | 2.73 | 65.92 | 0.00 | 0.00 | 29.15 | 0.04 | 0.83 |
| 02 | -1.63 | 0.97 | 5.01 | 33.09 | 0.00 | 52.33 | 0.97 | 1.59 |
| 04 | -0.63 | -1.11 | 4.62 | 62.70 | 7.55 | 18.12 | 2.09 | 1.45 |
| 07 | 0.05 | 1.93 | 26.86 | 6.72 | 0.00 | 33.50 | 0.26 | 0.83 |
| 09 | -2.92 | 0.09 | 0.00 | 27.10 | 0.00 | 65.73 | 0.88 | 6.03 |
| 12 | -2.47 | -1.07 | 0.00 | 51.76 | 2.32 | 40.42 | 2.10 | 1.30 |
| 13 | -2.88 | 0.13 | 0.00 | 21.77 | 00.00 | 66.17 | 0.18 | 6.35 |
| 17 | 2.28 | -0.97 | 40.95 | 22.40 | 16.93 | 2.36 | 0.00 | 0.73 |
| 22 | -1.44 | 1.12 | 7.36 | 38.51 | 00.00 | 51.55 | 0.03 | 1.98 |
| 23 | -1.09 | -2.63 | 0.20 | 14.79 | 31.11 | 21.8 | 0.10 | 3.31 |
| 31 | 3.02 | -0.37 | 68.10 | 4.29 | 14.67 | 2.09 | 4.61 | 1.98 |
| 32 | 3.34 | 0.99 | 92.81 | 3.44 | 1.14 | 0.31 | 0.00 | 4.00 |
| 33 | 3.14 | 0.52 | 76.10 | 6.23 | 2.22 | 0.40 | 0.00 | 2.09 |
| 36 | 3.23 | -0.68 | 63.25 | 10.90 | 11.82 | 0.41 | 4.33 | 1.06 |

Table S2: Summary statistics of the day in which data were collected for GPS tags applied to 43 female Tree Swallows on the twelve different farms (Appendix S2 Table S2).

| Farm ID | Nest box ID | Ordinal date | Mean distance traveled (SD) | | | Mean distance from nest box (SD) | | | Mean hourly temperature (SD) | | | Brood age | Brood size |
| --- | --- | --- | --- | --- | --- | --- | --- | --- | --- | --- | --- | --- | --- |
| 01 | 02 | 184 | 1471 | ± | 1324 | 368 | ± | 331 | 25 | ± | 6 | 10 | 5 |
| 01 | 08 | 167 | 1258 | ± | 1455 | 315 | ± | 363 | 19 | ± | 3 | 5 | 5 |
| 03 | 03 | 167 | 3202 | ± | 3364 | 800 | ± | 841 | 19 | ± | 4 | 7 | 5 |
| 03 | 10 | 163 | 483 | ± | 316 | 121 | ± | 79 | 19 | ± | 5 | 5 | 6 |
| 04 | 02 | 165 | 1231 | ± | 1025 | 308 | ± | 256 | 17 | ± | 2 | 5 | 7 |
| 04 | 04 | 165 | 904 | ± | 508 | 230 | ± | 126 | 17 | ± | 2 | 5 | 5 |
| 04 | 06 | 167 | 1228 | ± | 769 | 307 | ± | 192 | 20 | ± | 4 | 5 | 6 |
| 04 | 07 | 162 | 767 | ± | 585 | 192 | ± | 146 | 17 | ± | 1 | 6 | 6 |
| 04 | 08 | 163 | 1115 | ± | 1051 | 282 | ± | 261 | 21 | ± | 7 | 5 | 6 |
| 07 | 04 | 190 | 2968 | ± | 2100 | 742 | ± | 525 | 23 | ± | 4 | 6 | 5 |
| 07 | 09 | 184 | 1593 | ± | 2257 | 398 | ± | 564 | 22 | ± | 3 | 2 | 3 |
| 09 | 01 | 167 | 1383 | ± | 836 | 346 | ± | 209 | 19 | ± | 4 | 6 | 4 |
| 09 | 02 | 169 | 1705 | ± | 933 | 440 | ± | 254 | 22 | ± | 8 | 6 | 6 |
| 09 | 06 | 167 | 746 | ± | 902 | 186 | ± | 226 | 19 | ± | 4 | 6 | 1 |
| 09 | 07 | 164 | 1658 | ± | 3585 | 414 | ± | 896 | 15 | ± | 2 | 5 | 5 |
| 12 | 03 | 169 | 3718 | ± | 4784 | 929 | ± | 1196 | 22 | ± | 6 | 5 | 5 |
| 12 | 05 | 169 | 3461 | ± | 5664 | 865 | ± | 1416 | 22 | ± | 6 | 5 | 5 |
| 12 | 10 | 173 | 2109 | ± | 1007 | 527 | ± | 252 | 20 | ± | 5 | 7 | 5 |
| 13 | 02 | 186 | 3331 | ± | 2391 | 832 | ± | 598 | 27 | ± | 5 | 6 | 4 |
| 13 | 03 | 161 | 363 | ± | 210 | 91 | ± | 53 | 25 | ± | 5 | 5 | 4 |
| 13 | 04 | 167 | 2286 | ± | 2585 | 571 | ± | 646 | 19 | ± | 4 | 5 | 5 |
| 13 | 07 | 175 | 1556 | ± | 1320 | 389 | ± | 330 | 22 | ± | 7 | 7 | 5 |
| 17 | 05 | 174 | 3951 | ± | 3260 | 987 | ± | 815 | 25 | ± | 5 | 8 | 4 |
| 17 | 07 | 174 | 1996 | ± | 1579 | 499 | ± | 395 | 25 | ± | 5 | 6 | 2 |
| 22 | 01 | 173 | 4547 | 7 | 4889 | 1137 | ± | 1222 | 19 | ± | 3 | 5 | 6 |
| 22 | 02 | 179 | 3776 | ± | 3124 | 944 | ± | 781 | 25 | ± | 4 | 9 | 5 |
| 23 | 03 | 171 | 1067 | ± | 864 | 269 | ± | 214 | 19 | ± | 2 | 5 | 5 |
| 23 | 05 | 173 | 1376 | ± | 1450 | 344 | ± | 362 | 20 | ± | 3 | 5 | 5 |
| 23 | 07 | 173 | 889 | ± | 826 | 222 | ± | 207 | 20 | ± | 3 | 7 | 4 |
| 23 | 10 | 177 | 1636 | ± | 2391 | 465 | ± | 811 | 24 | ± | 4 | 9 | 7 |
| 31 | 04 | 166 | 596 | ± | 747 | 149 | ± | 187 | 18 | ± | 3 | 6 | 4 |
| 31 | 05 | 172 | 1883 | ± | 1059 | 470 | ± | 265 | 21 | ± | 4 | 10 | 5 |
| 31 | 07 | 168 | 1121 | ± | 602 | 284 | ± | 148 | 22 | ± | 4 | 6 | 5 |
| 31 | 08 | 166 | 628 | ± | 763 | 157 | ± | 191 | 18 | ± | 3 | 6 | 3 |
| 32 | 03 | 189 | 1272 | ± | 1108 | 323 | ± | 276 | 25 | ± | 7 | 7 | 3 |
| 32 | 08 | 176 | 2180 | ± | 2018 | 545 | ± | 505 | 21 | ± | 4 | 8 | 5 |
| 32 | 10 | 172 | 1073 | ± | 1604 | 268 | ± | 401 | 21 | ± | 4 | 6 | 5 |
| 33 | 07 | 176 | 2348 | ± | 2687 | 587 | ± | 672 | 19 | ± | 3 | 6 | 4 |
| 36 | 01 | 166 | 2206 | ± | 1638 | 551 | ± | 410 | 18 | ± | 3 | 6 | 3 |
| 36 | 02 | 166 | 3737 | ± | 3309 | 934 | ± | 827 | 18 | ± | 3 | 6 | 4 |
| 36 | 07 | 168 | 1311 | ± | 1145 | 328 | ± | 286 | 22 | ± | 4 | 6 | 3 |
| 36 | 08 | 168 | 2284 | ± | 1032 | 571 | ± | 258 | 22 | ± | 4 | 8 | 6 |
| 36 | 10 | 159 | 1114 | ± | 709 | 278 | ± | 177 | 21 | ± | 5 | 5 | 5 |

Table S3: Results of the model selection. Candidate models detail the covariate groups found within each model. Composition of each covariate group is detailed within Table 1. GLMMs with a negative binomial distribution with a log link function and whether a full hourly session was completed as a statistical offset were used to model the rate of food provisioning (N = 1,208 food provisioning sessions from 662 broods). GLMMs with a tweedie distribution and a log link function were used to model the biomass of food boluses (N = 741 food provisioning sessions 420 broods). The year, farm, and brood IDs were included as random effects in both sets of analyses. LMMs with the farm and individual id as random effects were used to model both the total hourly distance traveled and the mean hourly distance from a nest box (N = 529 GPS locations from 40 individuals). Interactions occurring with the Land covariate group occurred with the first component (Comp.1) as it explained the most variance within the higher-order habitat data.

| **Response** | **Candidate models** | **K** | ∆**AICc** | ***w*_i_** |
| --- | --- | --- | --- | --- |
| Bolus biomass | Base + Land | 21 | 0.00 | 0.24 |
|  | Base + Food + Land | 22 | 0.62 | 0.18 |
|  | Base + Land + Food * LSW | 23 | 1.28 | 0.13 |
|  | Base + Land * Demand | 24 | 1.46 | 0.12 |
|  | Base + Food + Land * Demand | 25 | 2.18 | 0.08 |
|  | Base | 18 | 2.30 | 0.08 |
|  | Base + Food + Land * LSW | 23 | 2.75 | 0.06 |
|  | Base + Food * Land | 23 | 2.75 | 0.06 |
|  | Base + Food | 19 | 4.10 | 0.03 |
|  | Base + Land + Food * Demand | 25 | 5.92 | 0.01 |
|  | Base + Food * Demand | 22 | 9.58 | 0.00 |
|  | Null | 6 | 44.69 | 0.00 |
| Bolus delivery | Base + Land * Demand | 23 | 0.00 | 0.29 |
|  | Base + Land | 20 | 0.33 | 0.25 |
|  | Base + Land + Food * LSW | 22 | 2.02 | 0.11 |
|  | Base + Food + Land * Demand | 24 | 2.03 | 0.10 |
|  | Base + Food + Land | 21 | 2.31 | 0.09 |
|  | Base | 17 | 3.26 | 0.06 |
|  | Base + Food * Land | 22 | 4.03 | 0.04 |
|  | Base + Food + Land * LSW | 22 | 4.36 | 0.03 |
|  | Base + Food | 18 | 4.93 | 0.02 |
|  | Base + Land + Food * Demand | 24 | 7.20 | 0.01 |
|  | Base + Food * Demand | 21 | 9.86 | 0.00 |
|  | Null | 5 | 54.20 | 0.00 |
| Total distance | Base + Land*WT | 18 | 0.00 | 0.76 |
|  | Base + Land | 17 | 3.22 | 0.15 |
|  | Base + Land*Food | 19 | 5.67 | 0.04 |
|  | Base + Land*Demand | 20 | 6.46 | 0.03 |
|  | Base | 14 | 8.29 | 0.01 |
|  | Null | 4 | 97.77 | 0.00 |
| Distance from box | Base + Land | 17 | 0.00 | 0.36 |
|  | Base + Land*WT | 18 | 0.39 | 0.30 |
|  | Base + Land*Food | 19 | 2.02 | 0.13 |
|  | Base | 14 | 2.17 | 0.12 |
|  | Base + Land*Demand | 20 | 2.68 | 0.09 |
|  | Null | 4 | 64.35 | 0.00 |

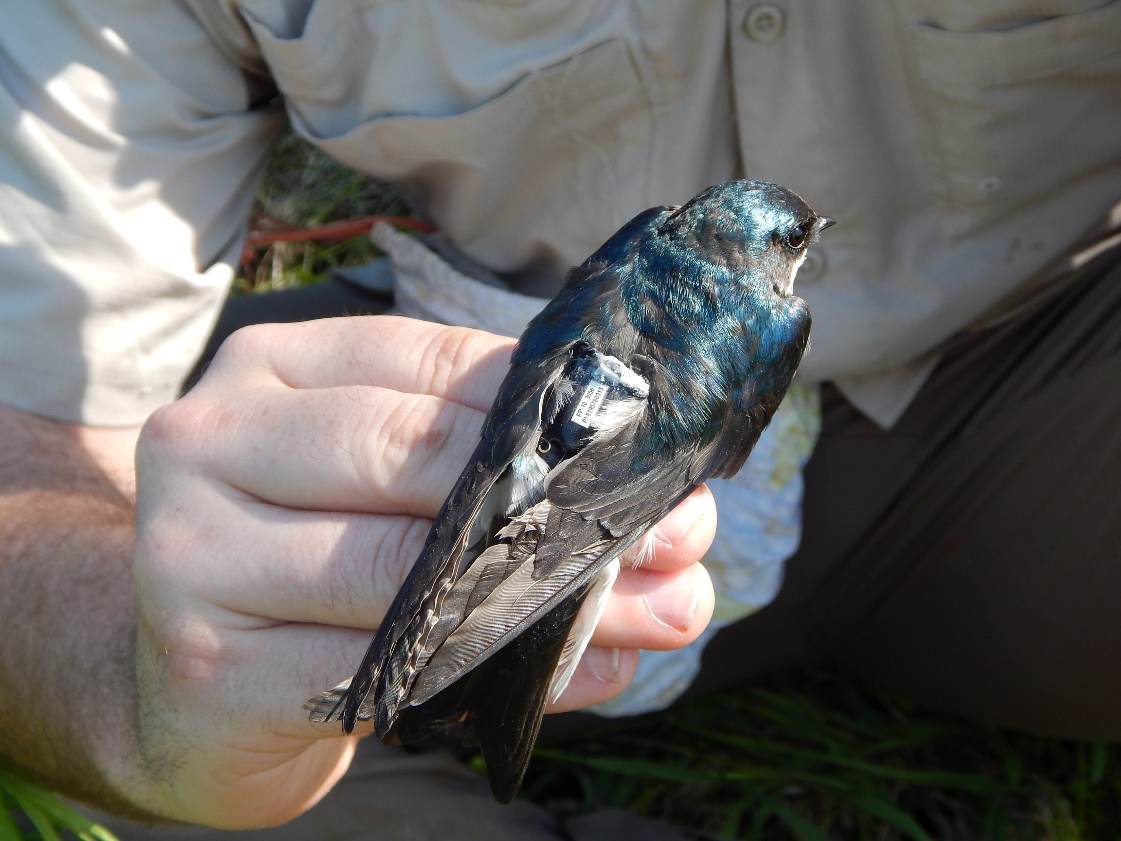

Figure S1: A PinPoint GPS tag attached to a breeding female Tree Swallow. Tags weighed between 0.9 g and 1.2 g, including the harness, and were fitted over the synsacrum using leg loops (Rappole and Tipton 1991). Once the harness was fitted, the loose string attached on top of the tag using Cyanoacrylate glue (white material at proximal end of tag) was cut.

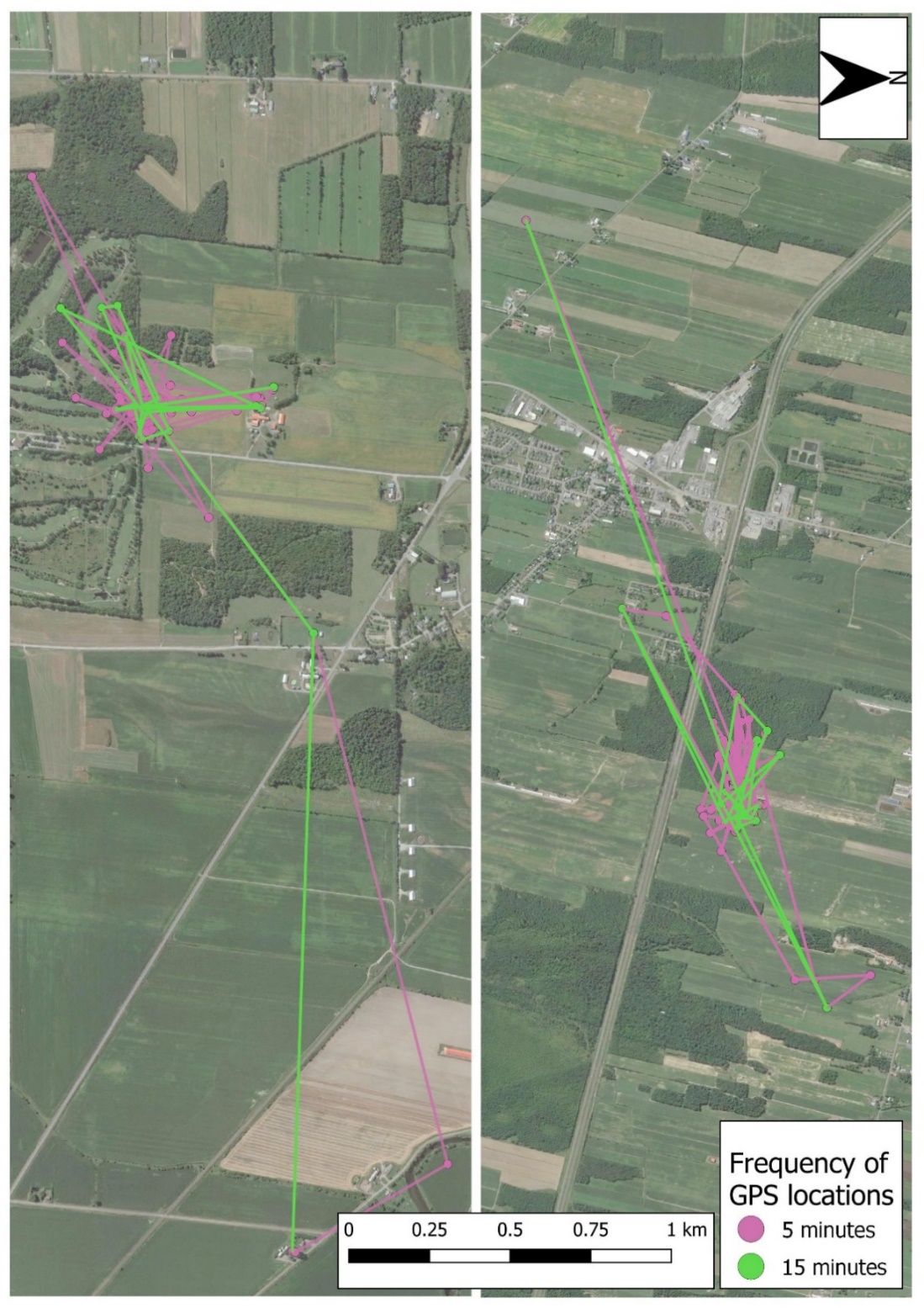

Figure S2: Map of GPS location and associated paths for two females in which GPS locations were recorded every 5 minutes compared to every 15 minutes. In pink are 5-minute locations and green 15-minute locations. Despite the collection of more spatial data, no over arching trends appear to be missed by focusing sampling on every 15 minutes.

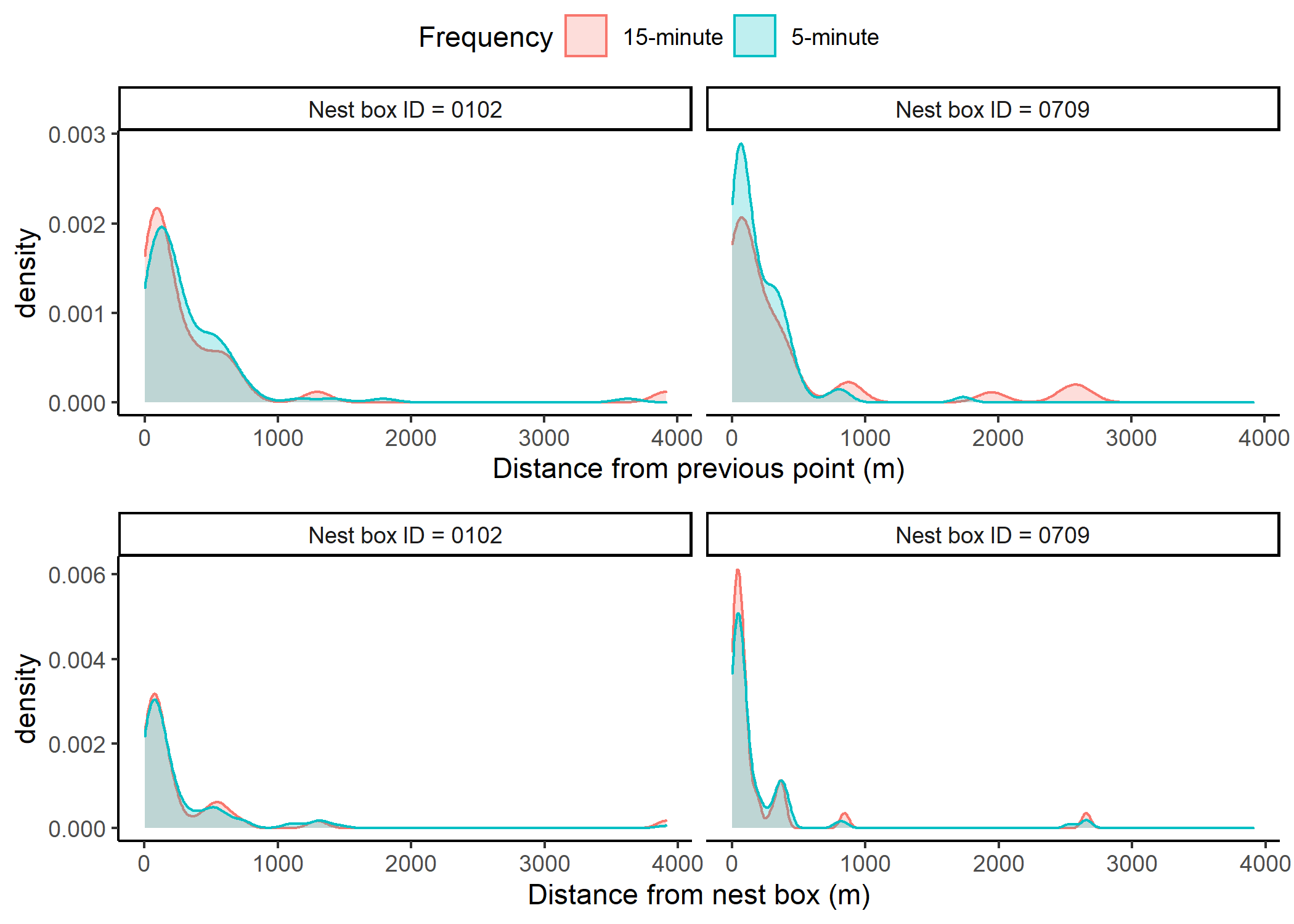

Figure S3: Comparison of kernel densities of the distances between GPS locations and both the previous GPS location and the individual’s nest box for two breeding females. Comparison is for GPS locations recorded every 5 minutes and every 15 minutes.

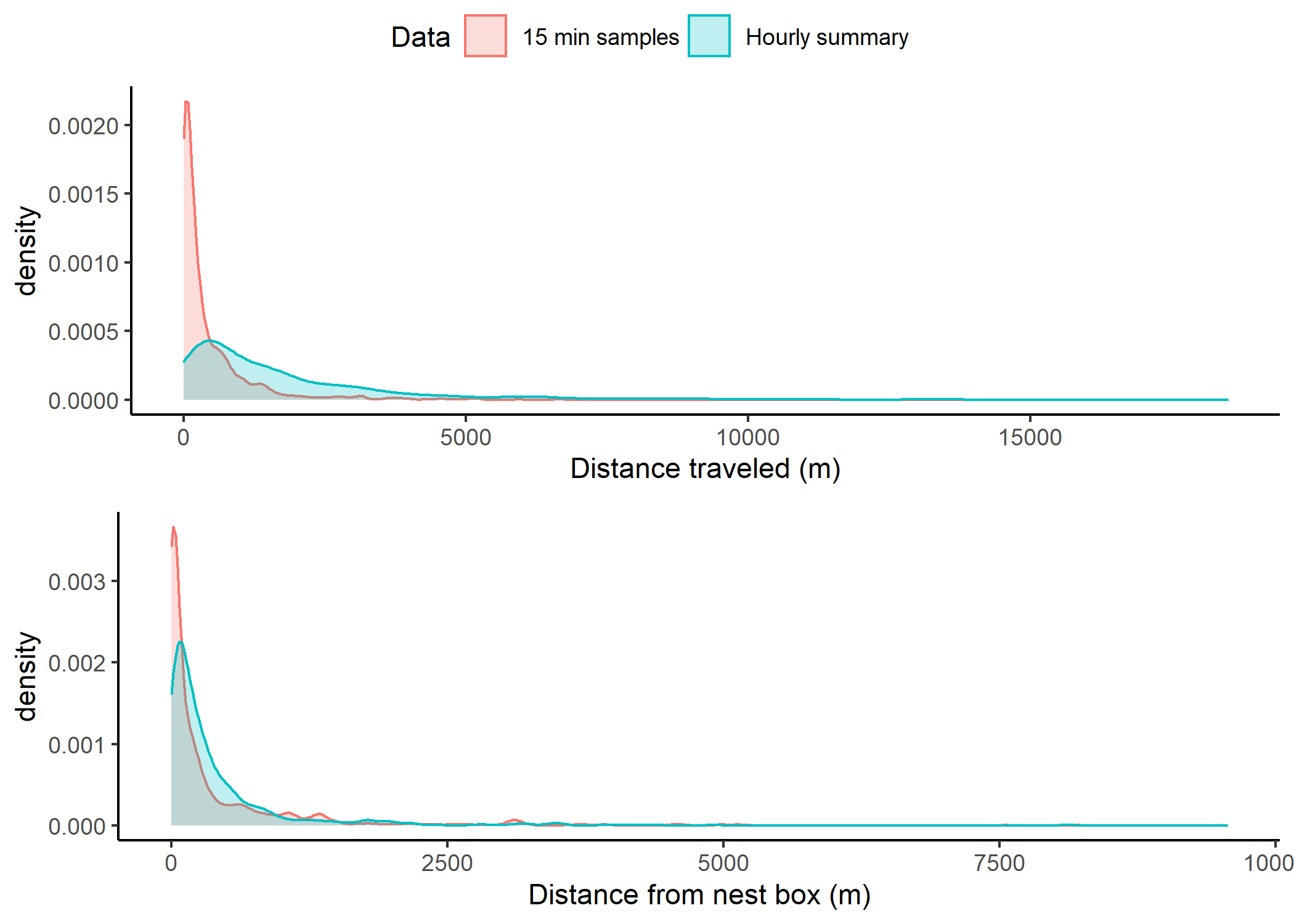

Figure S4: Comparison of kernel densities of the distances between GPS locations and nest boxes as well as between locations defined by sampling events or summarized during hourly summaries. Distance traveled for 15-minute intervals is the distance between a GPS location and the previous location, while for hourly summary is the summation of distance traveled during one-hour sampling events. Distance from nest box for 15-minute intervals refers to the distance a GPS location was from the bird’s nest box, while for hourly summary is the mean of this value during one-hour sampling event.

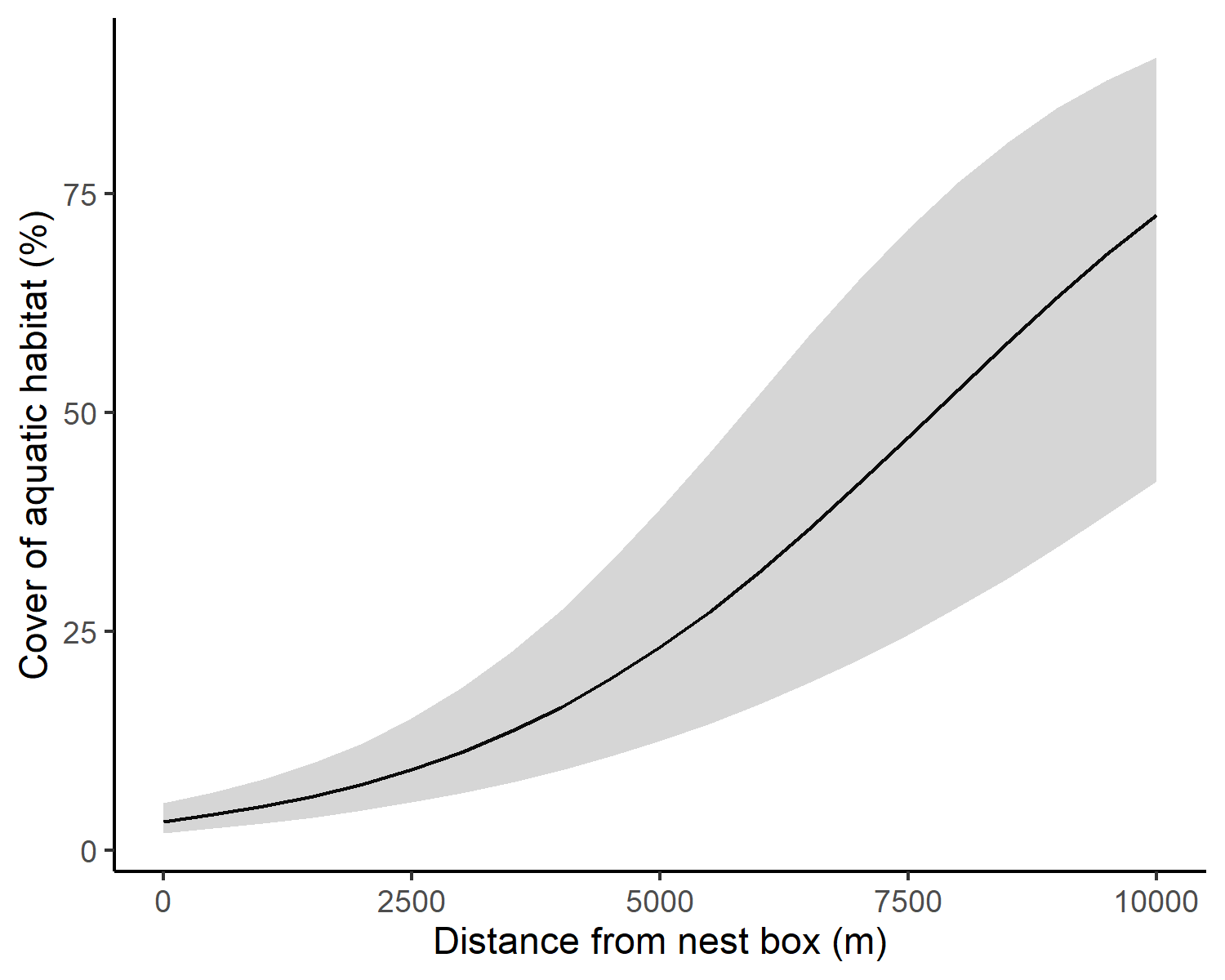

Figure S5: Percent cover of aquatic habitat within 100 m of food provisioning female Tree Swallows as a function of the distance the females were from their respective nest box. Presented are predictions and 95% confidence intervals. Percent cover was modeled using GLMMs with a beta-binomial distribution and a logit link function. Models included farm and nest box ID as random factors to account for the hierarchical nature of the data (N = 2,516 GPS locations across 43 individuals).

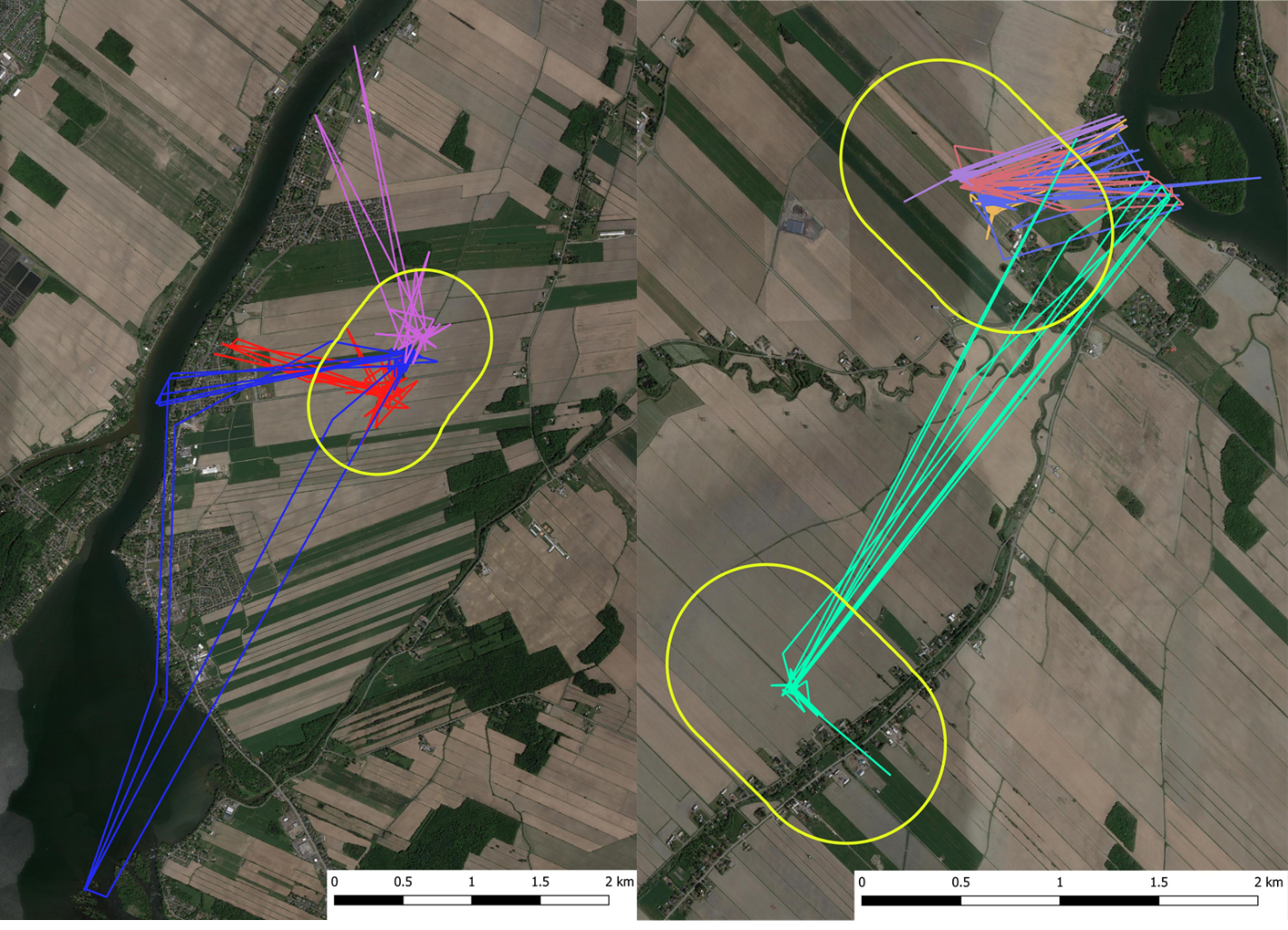

Figure S6: Aerial image of agricultural landscapes and GPS locations recorded for three farms. Color of lines indicate separate individuals. Yellow polygons depict 500-m buffers around each nest box of a farm that have been coalesced to create one buffer per farm. We preprogrammed tags so GPS fixes (“SWIFT” fixes, nominal horizontal accuracy ± 10 m) were taken every 15 min between 5:00 and 20:00 local time (N = 60 locations) the day after deployment, allowing for a habituation period. Figure is intended to depict traveling events towards water bodies.

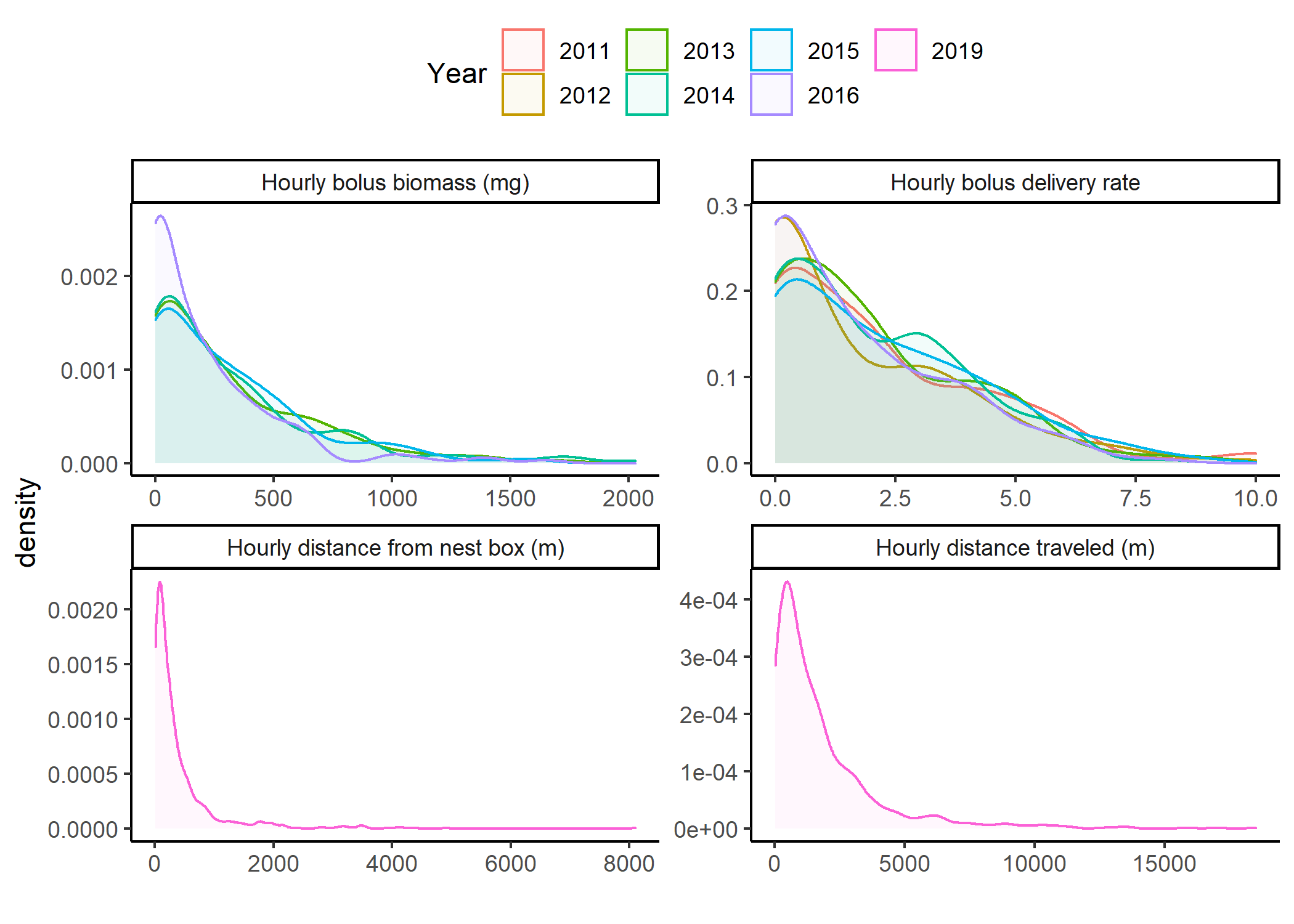

Figure S7: Kernel density plot of hourly response variables broken down by year of sampling.

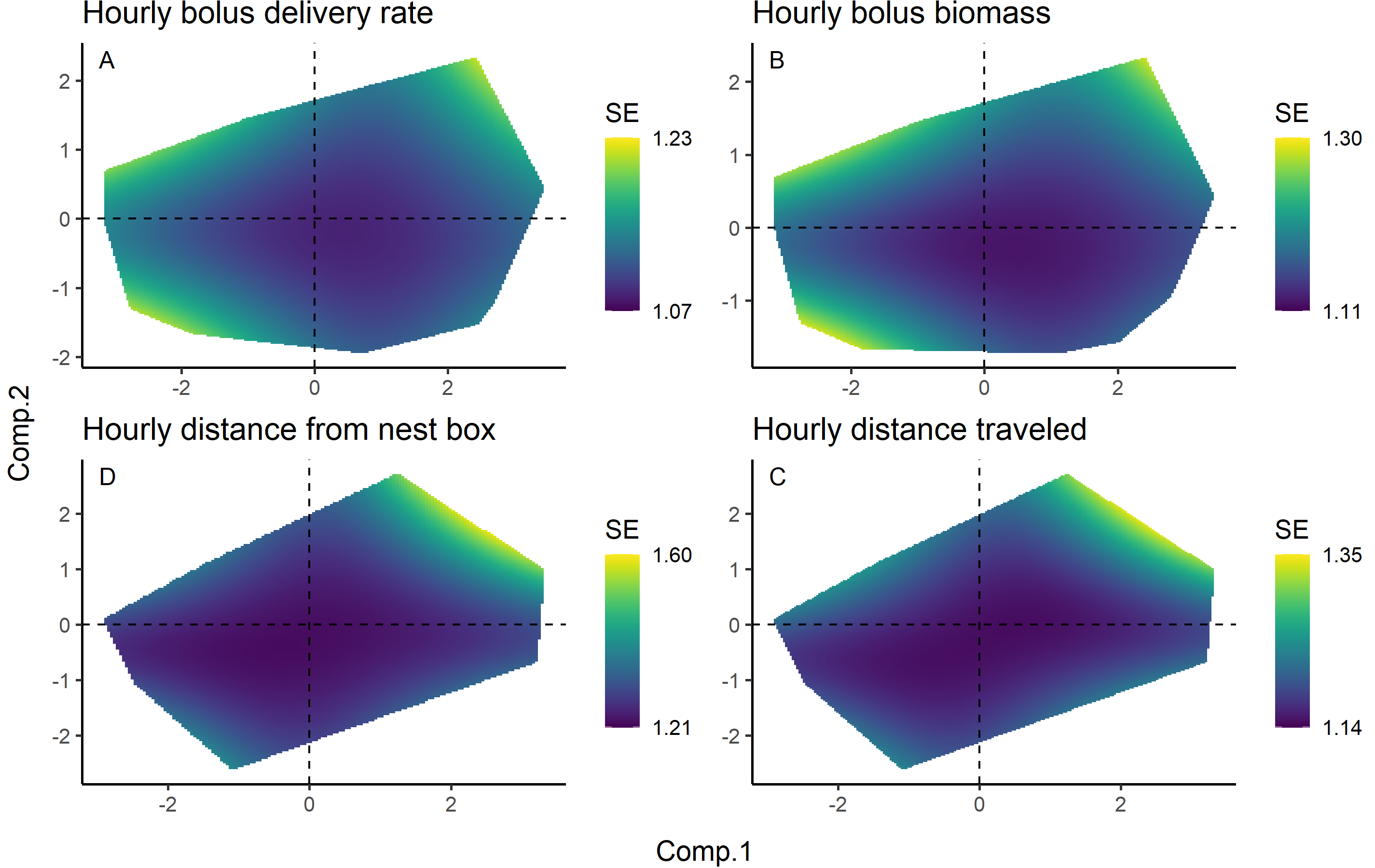

Figure S8: Prediction surface demonstrating the unconditional standard error of the predictions from Fig 4 for each response variable. Standard errors are on the response scale for each response variable. Surfaces are the two-dimensional unconditional standard error of each response variable over the observed range of values along the first and second component values of the PCA defining landscape context found in Fig 3 and averaged according to the model weights in Table S3.

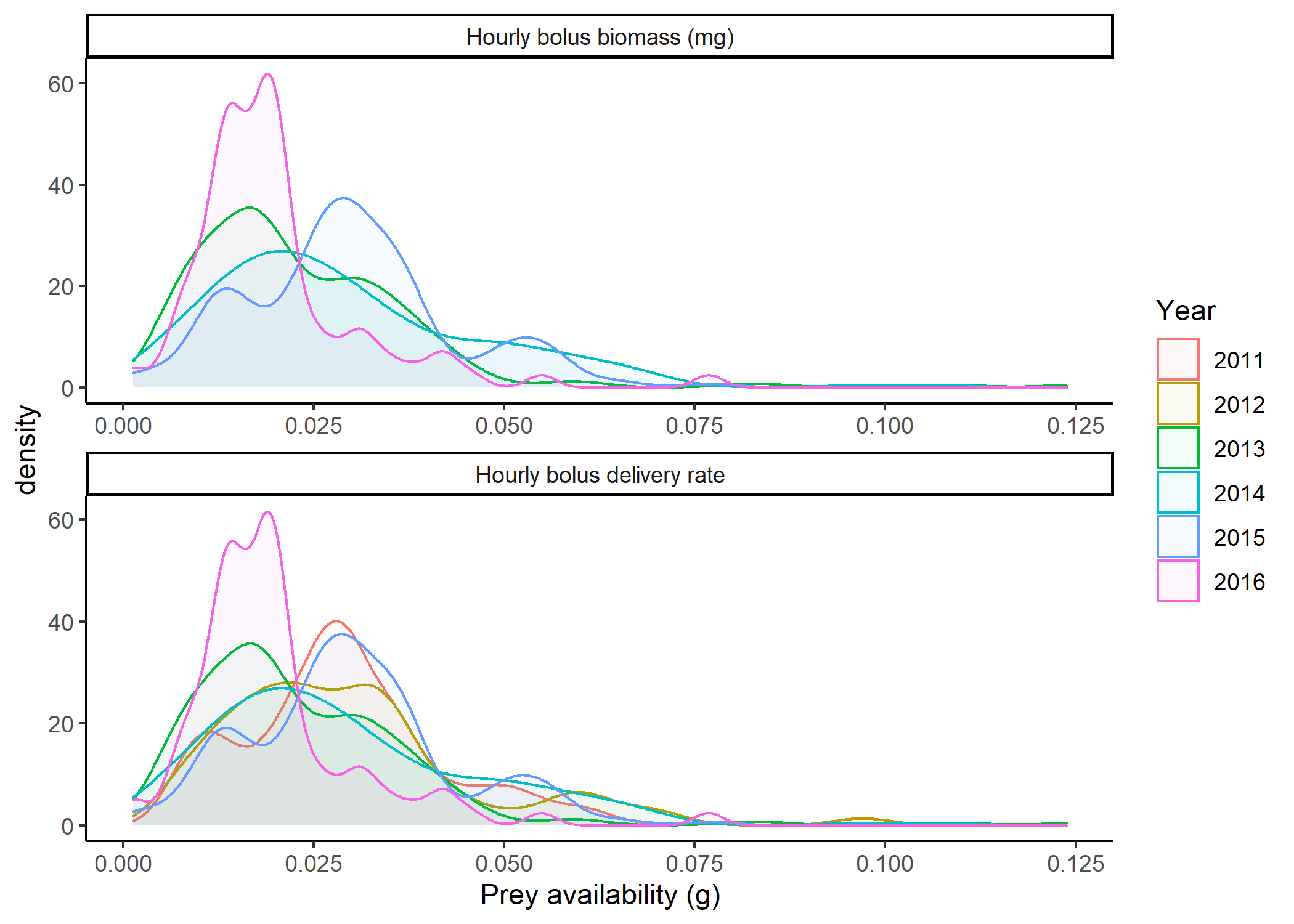

Figure S9: Kernel density plots of prey availability for each response variable broken down by year of sampling.

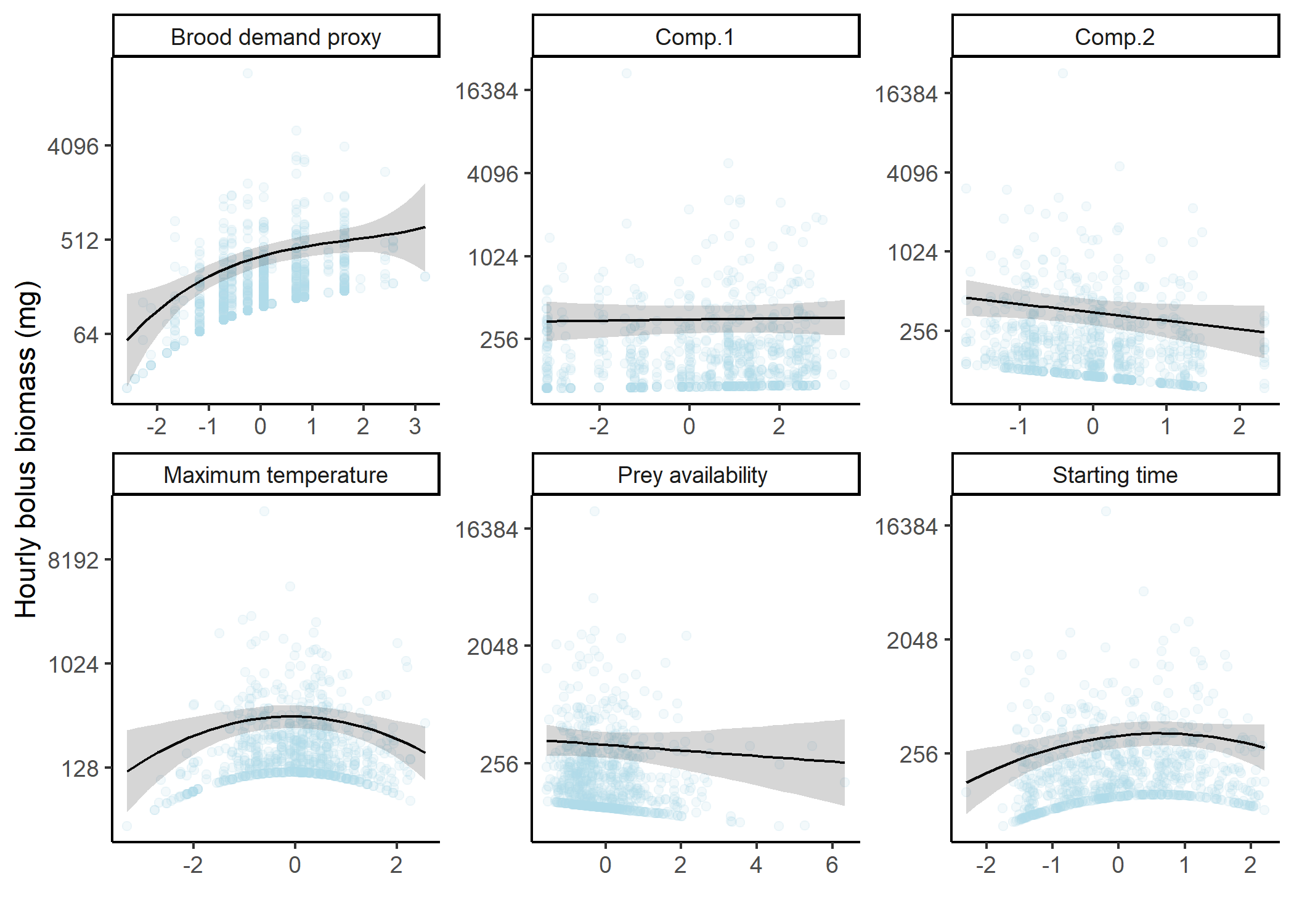

Figure S10: Model-averaged predictions and partial residuals of hourly biomass of food boluses with unconditional 95% confidence intervals against each covariate of interest. In each case, the values of all other covariates have been set to their mean and the predictor under consideration has been scaled using a z-transformation. See Table 1 for meaning of covariate acronyms and predictions are averaged according to the model weights in Table S3.

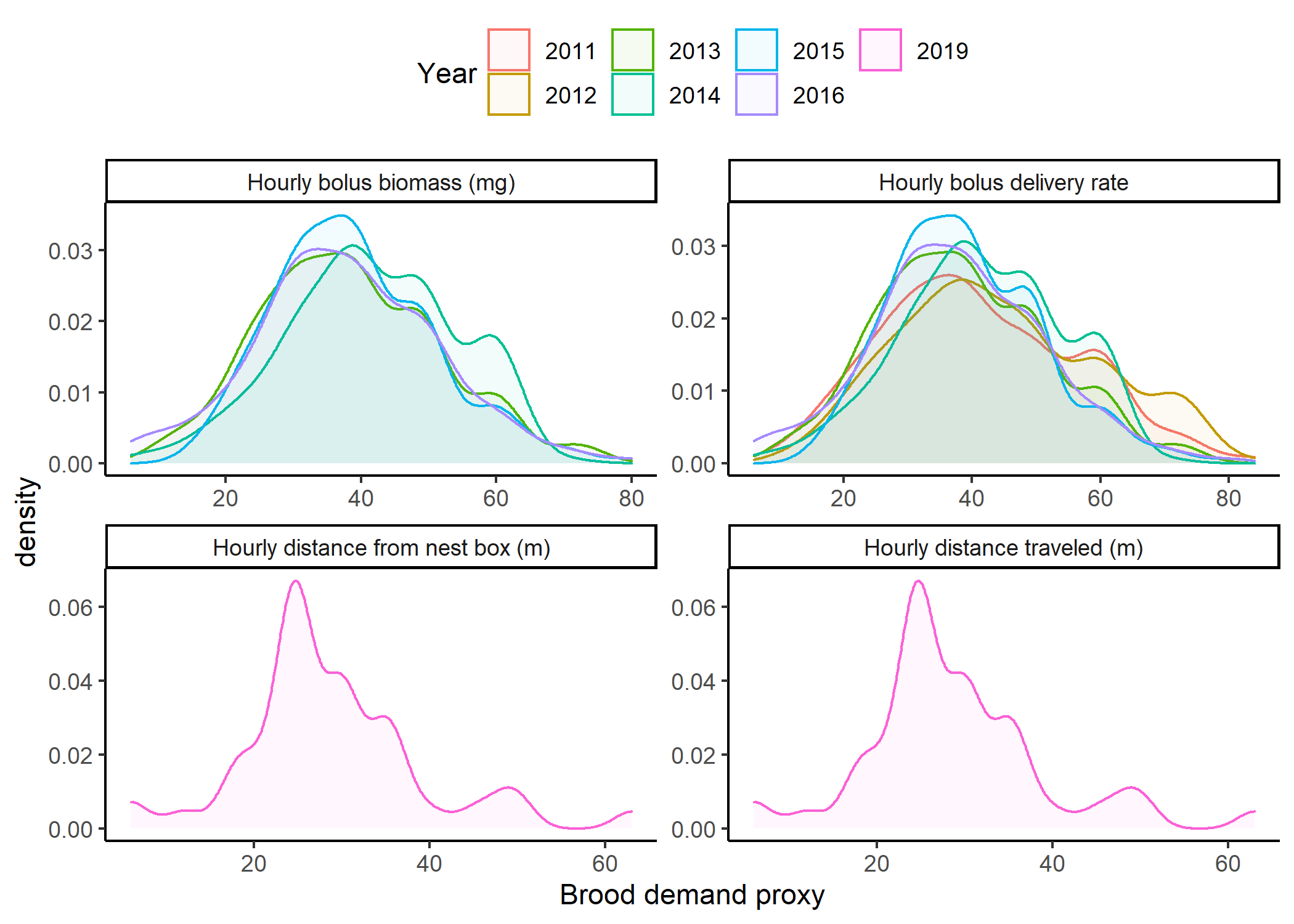

Figure S11: Kernel density plots of brood demand proxy for each response variable broken down by year of sampling. Proxy of brood demand was the product of the brood size and brood age at the time of sampling.

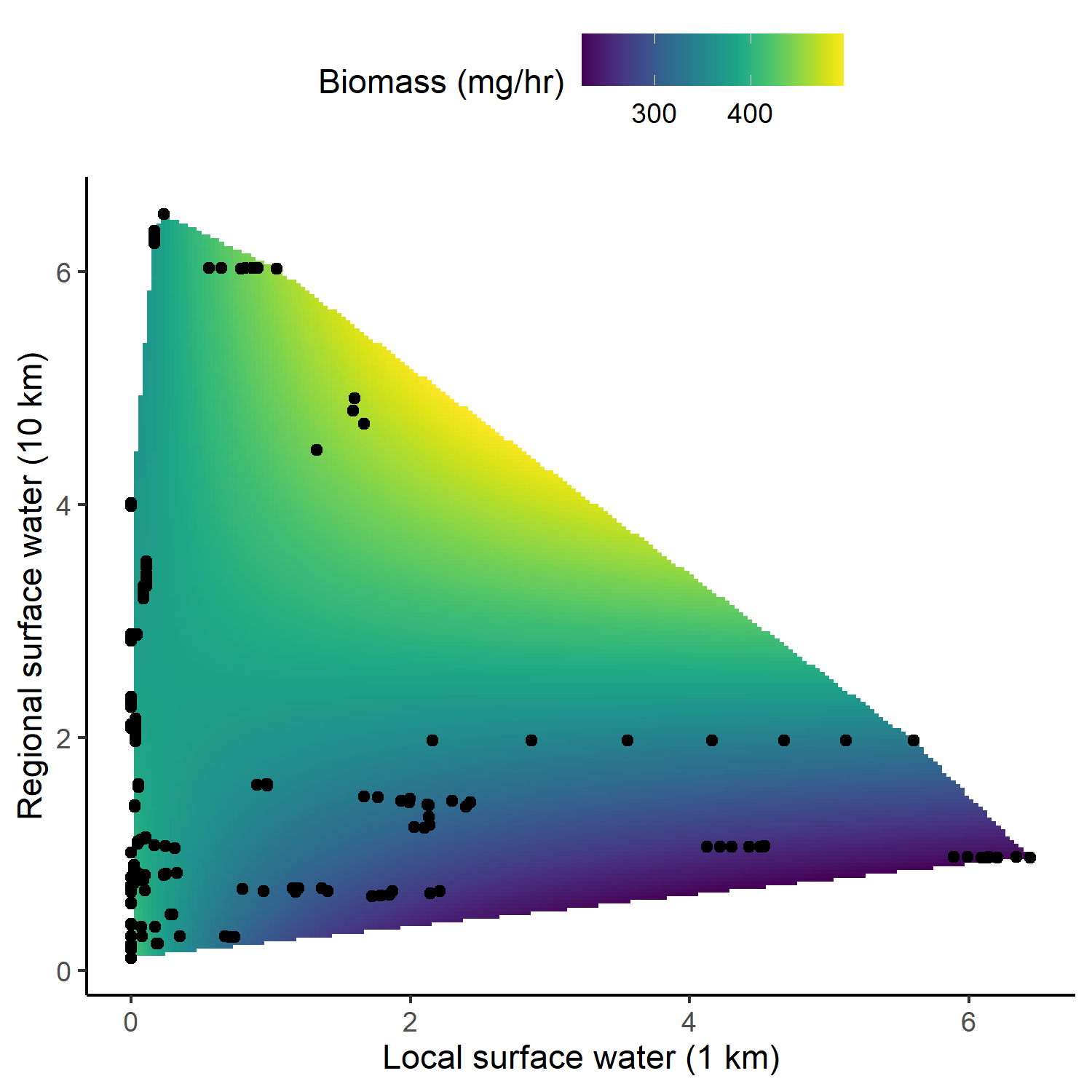

Figure S12: Model-averaged predictions for hourly biomass of food boluses against the percent cover of surface water within 1 km and 10 km around a nest box. around a nest box. In each case, the values of all other covariates have been set to their mean.

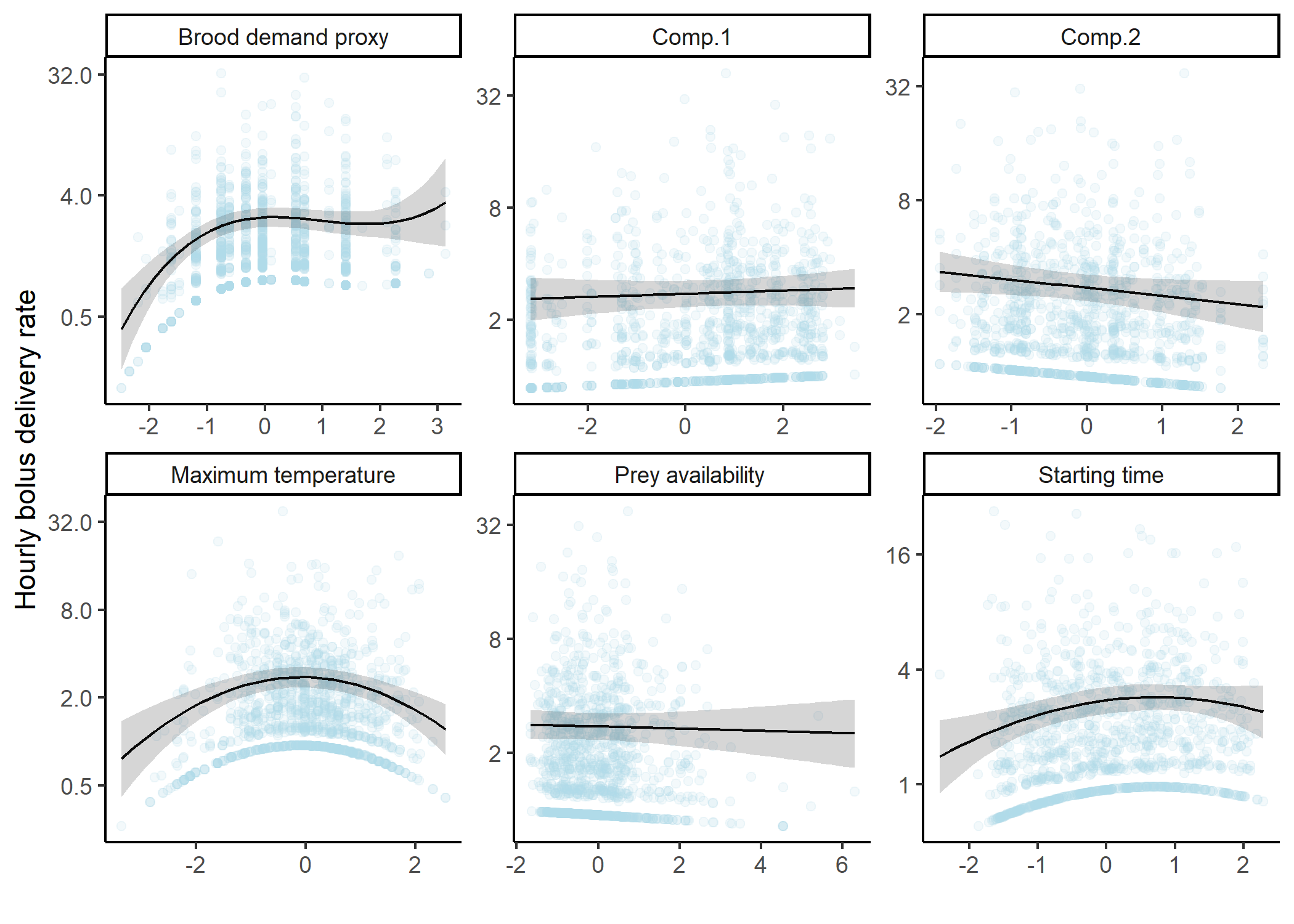

Figure S13: Model-averaged predictions and partial residuals of hourly number of food boluses delivered with unconditional 95% confidence intervals against each covariate of interest. In each case, the values of all other covariates have been set to their mean and the predictor under consideration has been scaled using a z-transformation. See Table 1 for meaning of covariate acronyms and predictions are averaged according to the model weights in Table S3.

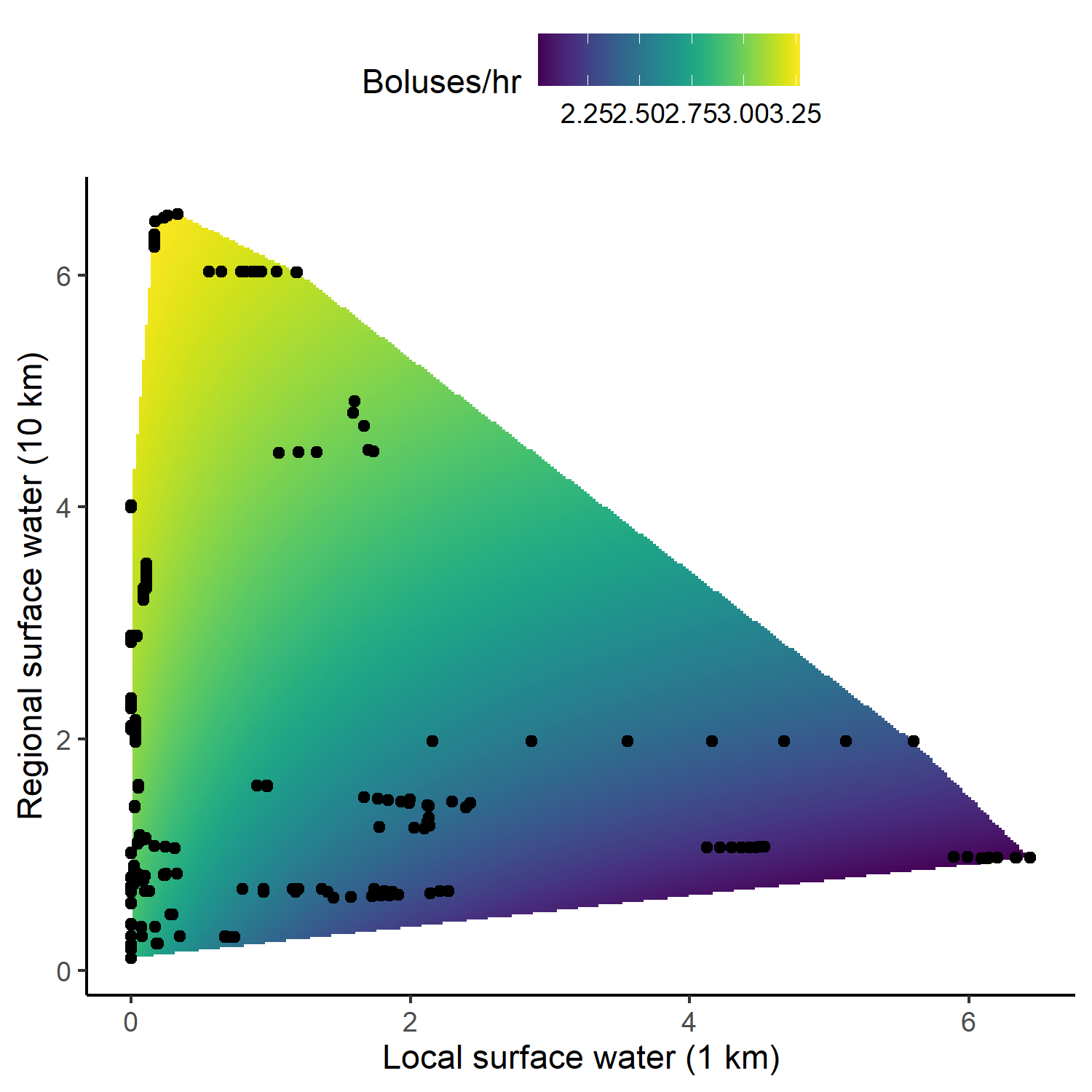

Figure S14: Model-averaged predictions of hourly number of boluses against the percent cover of surface water within 1 km and 10 km around a nest box. around a nest box. In each case, the values of all other covariates have been set to their mean.

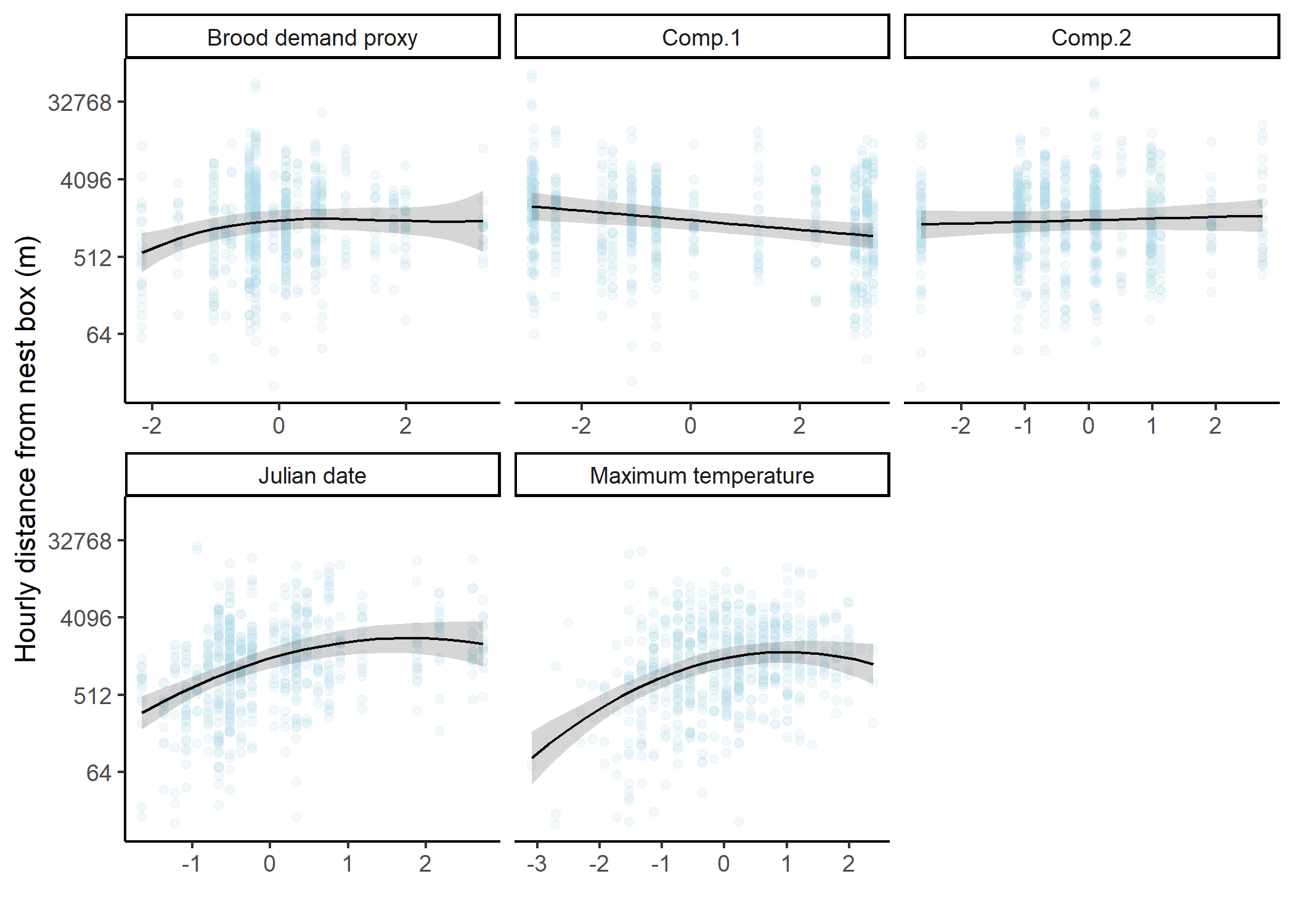

Figure S15: Model-averaged predictions and partial residuals of hourly total distance traveled (m) with unconditional 95% confidence intervals against each covariate of interest. In each case, the values of all other covariates have been set to their mean and the predictor under consideration has been scaled using a z-transformation. See Table 1 for meaning of covariate acronyms and predictions are averaged according to the model weights in Table S3.

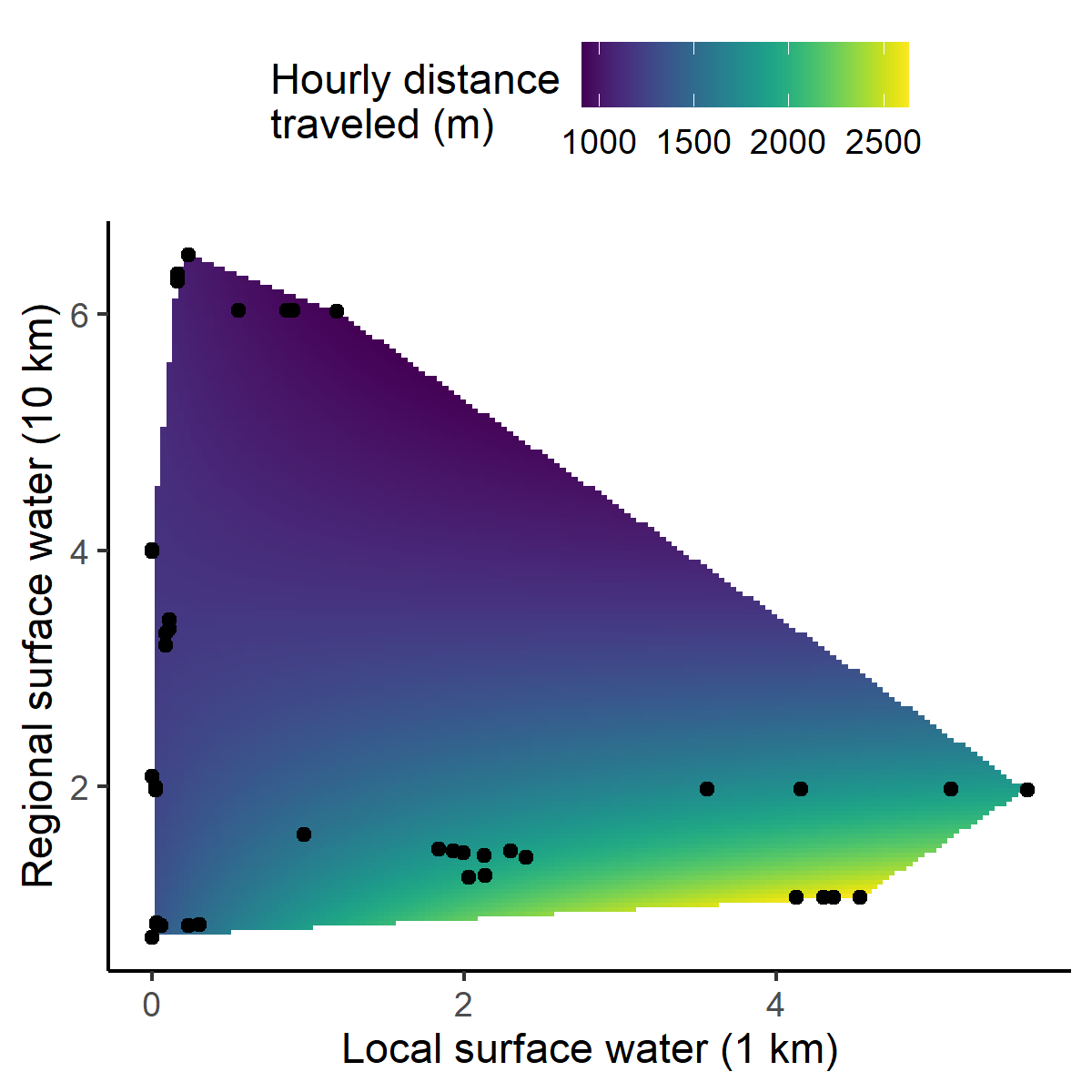

Figure S16: Model-averaged predictions of hourly distance traveled (m) against the percent cover of surface water within 1 km and 10 km around a nest box. The values of all other covariates have been set to their mean.

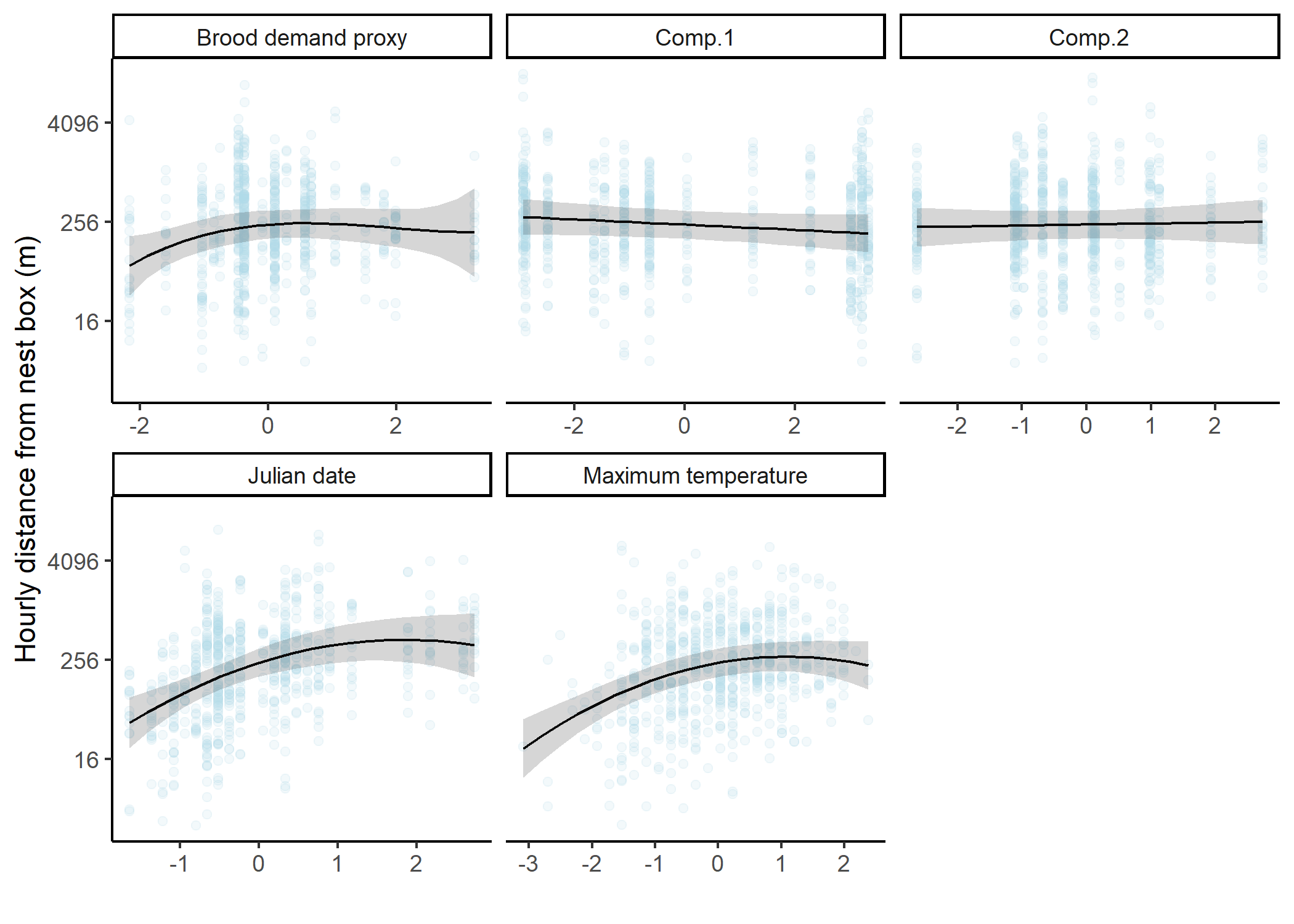

Figure S17: Model-averaged predictions and partial residuals of mean hourly distance from nest box (m) with unconditional 95% confidence intervals against each covariate of interest. In each case, the values of all other covariates have been set to their mean and the predictor under consideration has been scaled using a z-transformation. See Table 1 for meaning of covariate acronyms and predictions are averaged according to the model weights in Table S3.

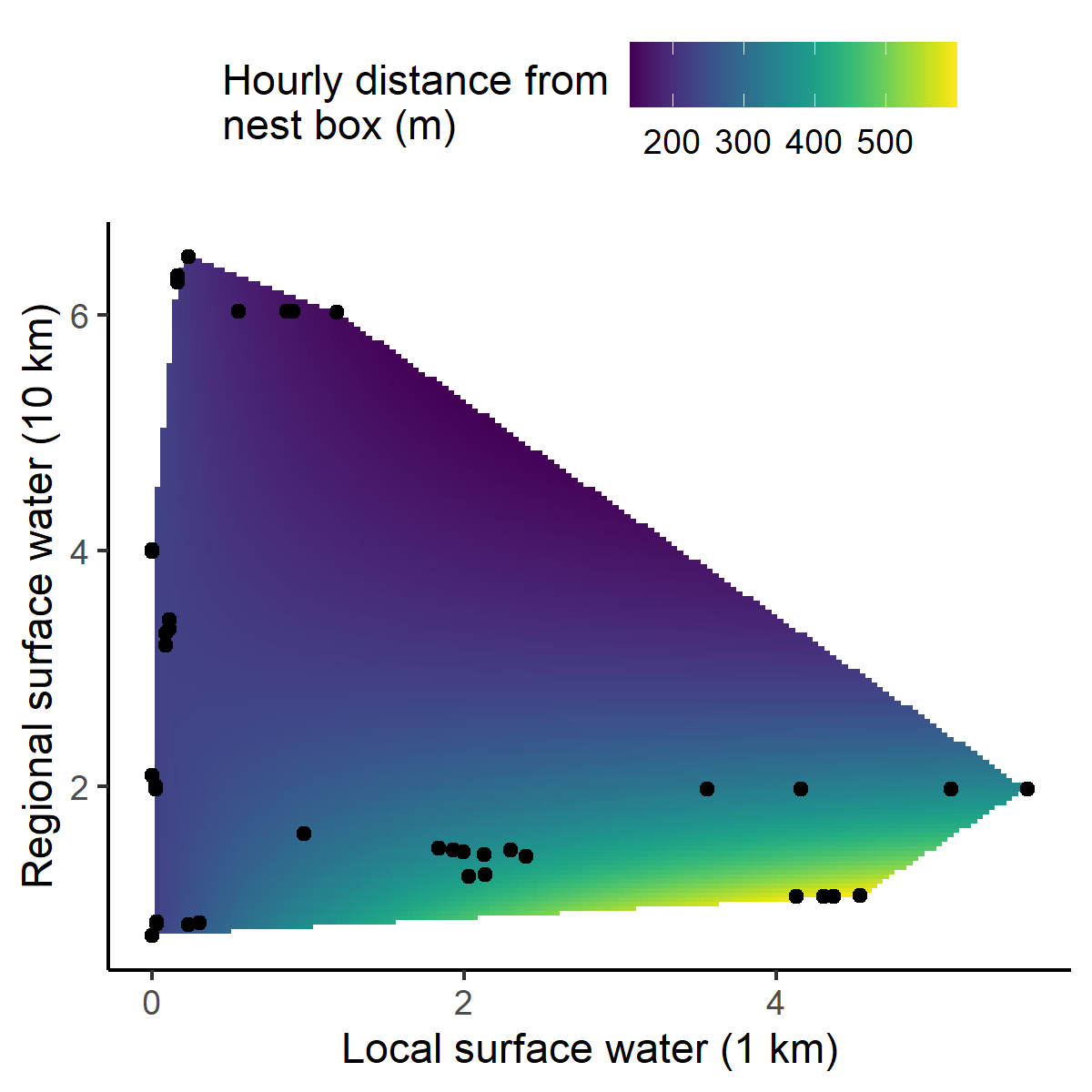

Figure S18: Model-averaged predictions of hourly distance from nest box (m) against the percent cover of surface water within 1 km and 10 km around a nest box. The values of all other covariates have been set to their mean.
