## AppendixS3 for "Negative effects of agricultural intensification on the food provisioning rate of a declining aerial insectivore"

### Appendix S3

##### **Determination of spatial scale for the calculation of aquatic habitat**

GPS locations greater than 0.5 km from nest boxes (19.3% of locations) frequently occurred over or close to aquatic bodies (e.g. rivers, wetlands and ponds), supporting the contention that these features are important to Tree Swallows (Elgin et al. 2020, Berzins et al. 2020). These habitats are however rare within our study area and were considered as potential confounding factors of the observed foraging responses. Moreover, we hypothesized that the influence of aquatic features located far from nests (Regional) may depend on their local availability (Local). We therefore undertook a modeling endeavor aimed at identifying the spatial extent and the type of aquatic features most influential to the number of delivered food boluses. We compared models that were identical in all respects with the exception of the spatial scale and type of aquatic habitat considered (model: Base + Land in Table S3). We explored two separate aquatic habitats: wetlands and open bodies of water. Wetland data were derived from available polygon layers produced by Ducks Unlimited that accurately maps wetlands greater than 0.5 hectare (Ducks unlimited, 2019). Open bodies of water were represented by polygon layers acquired from the Canadian National Hydro Network (NHN, 2020). The relative cover of aquatic habitats closes to nest boxes being significantly low, we focused on spatial scales spanning from 1 km to 10 km at 500 m increments. For each spatial scale, the percent cover of both aquatic habitats, plus their total non-overlapping cover were calculated using the sf (Pebesma 2018) R package. We then evaluated the second order Akaike information criterion (AICc) and the associated Akaike weight (*w*) of competing models. The spatial scales and aquatic habitat considered in all subsequent models of the present study would be chosen based on the model resulting in the greatest Akaike weight. Several models could not converge, and we found numerical stability was greatest when treating aquatic habitats as open bodies of water. Focusing on this aquatic habitat type, a local maximum of AICc weight (*w*) was observed when including the percent cover of surface water calculated within 1 km (local percent cover) and 10 km (regional percent cover) from the nest, including their interaction (Fig. S3). These model terms were included in all models for all foraging proxy.

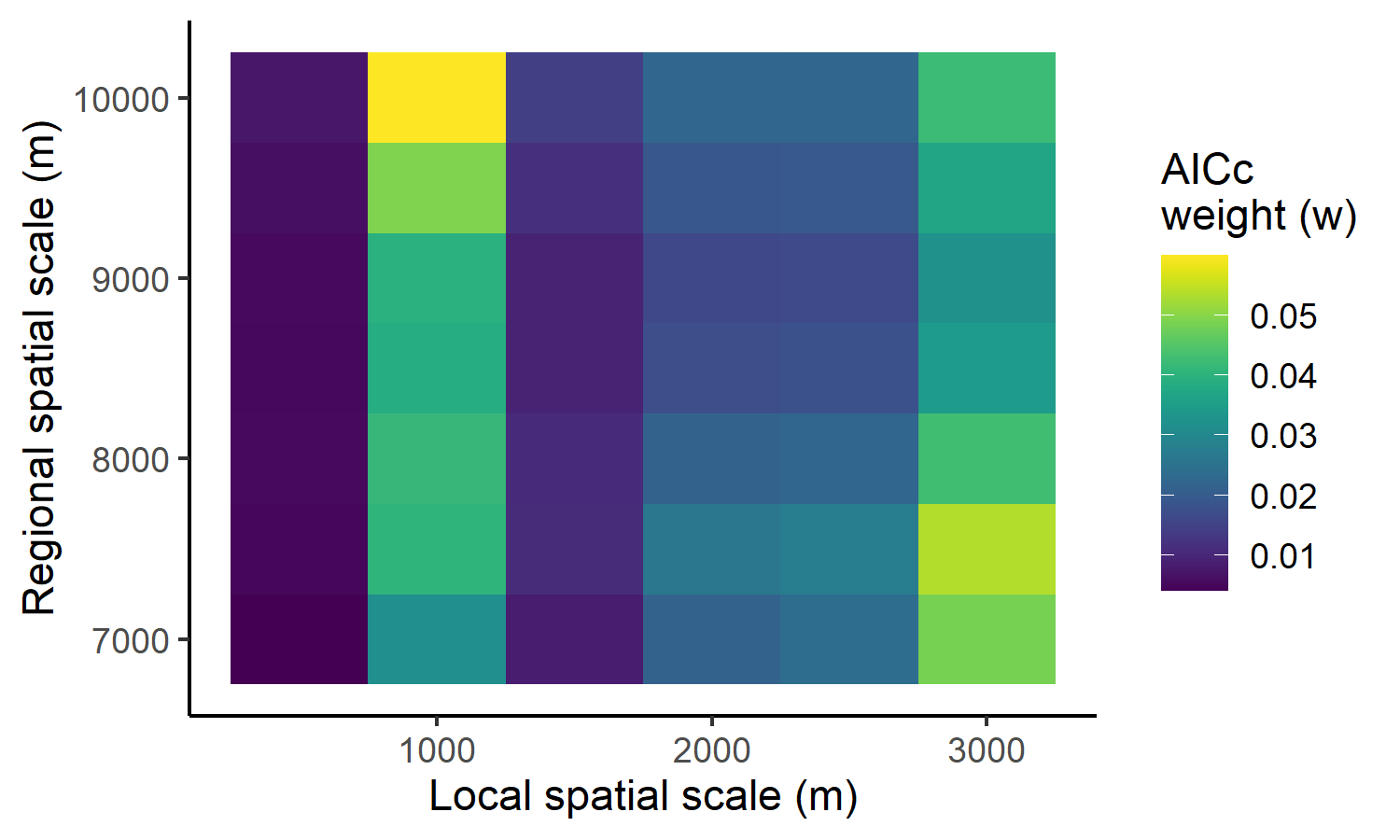

**Figure S1:**	Visual representation of model selection process for determining the spatial scale and aquatic habitat type most influential to hourly bolus delivery rate. Due to frequent non-convergence, analyses focused on aquatic habitats characterized as open bodies of water. The x and y axis are the local and regional spatial scale to which the percent cover of aquatic habitat was calculated, respectively. The model weight (*w*) is represented by depth of pixel color.
