## AppendixS4 for "Negative effects of agricultural intensification on the food provisioning rate of a declining aerial insectivore"

**Appendix S4**

#### **Analyses of individual bolus biomass**

The biomass of individual food boluses was a continuous right-skewed (conditional) response and was consequently modeled with GLMMs using a Gamma distribution and a log link function (Lo and Andrews 2015). We included year, as well as nest box and brood identification IDs, as random factors in order to account for the hierarchical structure of these data (i.e., broods nested within farms nested within years). We further accounted for factors potentially masking the main effects of interests. These factors were either predicted or presented to be influential to the foraging behavior of Tree Swallows (see main text). Factors included the time of day, maximum temperature, presence of precipitation, brood size, age (second-year (SY) vs. after-second-year (ASY)) of the breeding female, and the interaction between the local and regional percent cover of water. Residual diagnostics suggested a parabolic relationship with the time of day and the maximum temperature and were subsequently treated as second degree polynomials. We evaluated the relative effects of key individual explanatory variables by comparing the same set of competing models used for both the total hourly biomass and hourly number of boluses delivered (Table 1 and Table S2). The effects of key individual variables, including interactions, were estimated via multi-model inference whereby predictions were calculated by model-averaging with shrinkage and shown with their 95% unconditional confidence intervals (Burnham and Anderson 2002).

Table S1:	Results of the model selection for the biomass of individual boluses. Candidate models detail the covariate groups found within each model. Composition of each covariate group is detailed within Table 1. GLMMs with a Gamma distribution and a log link function were used to model these data (N = 815). The year, farm, and brood IDs were included as random effects in both sets of analyses. The session ID was included as another random effect in the analyses of bolus biomass to account for multiple boluses collected during each food provisioning session.

| **Response** | **Candidate models** | **K** | ∆**AICc** | ***w*_i_** |
| --- | --- | --- | --- | --- |
| Individual Bolus biomass | Base | 17 | 0.00 | 0.46 |
|  | Base + Food * Demand | 21 | 1.00 | 0.28 |
|  | Base + Food | 18 | 1.89 | 0.18 |
|  | Base + Land | 20 | 5.96 | 0.02 |
|  | Base + Land + Food * Demand | 24 | 6.83 | 0.02 |
|  | Base + Food + Land | 21 | 7.88 | 0.01 |
|  | Base + Land * Demand | 23 | 8.07 | 0.01 |
|  | Base + Food * Land | 22 | 8.98 | 0.01 |
|  | Base + Land + Food * LSW | 22 | 9.11 | 0.00 |
|  | Base + Food + Land * LSW | 22 | 9.75 | 0.00 |
|  | Base + Food + Land * Demand | 24 | 9.97 | 0.00 |
|  | Null | 5 | 68.32 | 0.00 |

Table S2:	Standardized coefficient estimates, and their 95% confidence intervals included in the top model predicting the biomass of individual boluses. See Table 1 for definitions of covariates and Table S2 for the outcome of model selection. Coefficients in bold indicate confidence intervals not overlapping zero. Numbers next to acronym refer to the order of the polynomial term.

| **Covariate** | **Estimate** | **Lower 95%** | **Upper 95%** |
| --- | --- | --- | --- |
| Female age (SY) | 0.106 | 0.075 | -0.041 |
| Rain (Yes) | -0.024 | 0.098 | -0.216 |
| **DM1** | **7.805** | **0.944** | **5.954** |
| DM2 | -0.651 | 0.898 | -2.412 |
| DM3 | -3.062 | 0.846 | -4.720 |
| ST1 | -0.145 | 0.992 | -2.088 |
| ST2 | -0.377 | 0.950 | -2.239 |
| TMax1 | -2.425 | 0.950 | -4.287 |
| TMax2 | 0.433 | 0.866 | -1.265 |
| LSW | -0.037 | 0.027 | -0.090 |
| LSW:RSW | 0.057 | 0.043 | -0.027 |
| RSW | -0.004 | 0.028 | -0.059 |
